# Identification of Key Pathways and Genes in Dementia via Integrated Bioinformatics Analysis

**DOI:** 10.1101/2021.04.18.440371

**Authors:** Basavaraj Vastrad, Chanabasayya Vastrad

## Abstract

To provide a better understanding of dementia at the molecular level, this study aimed to identify the genes and key pathways associated with dementia by using integrated bioinformatics analysis. Based on the expression profiling by high throughput sequencing dataset GSE153960 derived from the Gene Expression Omnibus (GEO), the differentially expressed genes (DEGs) between patients with dementia and healthy controls were identified. With DEGs, we performed a series of functional enrichment analyses. Then, a protein–protein interaction (PPI) network, modules, miRNA-hub gene regulatory network and TF-hub gene regulatory network was constructed, analyzed and visualized, with which the hub genes miRNAs and TFs nodes were screened out. Finally, validation of hub genes was performed by using receiver operating characteristic curve (ROC) analysis. A total of 948 DEGs were screened out, among which 475 genes were up regulated; while 473 were down regulated. Functional enrichment analyses indicated that DEGs were mainly involved in defense response, ion transport, neutrophil degranulation and neuronal system. The hub genes (CDK1, TOP2A, MAD2L1, RSL24D1, CDKN1A, NOTCH3, MYB, PWP2, WNT7B and HSPA12B) were identified from PPI network, modules, miRNA-hub gene regulatory network and TF-hub gene regulatory network. We identified a series of key genes along with the pathways that were most closely related with dementia initiation and progression. Our results provide a more detailed molecular mechanism for the advancement of dementia, shedding light on the potential biomarkers and therapeutic targets.

## Introduction

Dementia is a neuropsychiatric syndrome characterized by losses of cognitive and emotional abilities [Cantone et al. 2014]. Dementia patients show varying levels of behavior disturbance includes agitation, aggression, and rejection of care [Marx et al. 2019]. The numbers of cases of dementia are increasing globally and it has become an essential mental health concern. According to a survey, the prevalence of dementia is expected to reaching 110 million in 2050 [Lyketsos, 2020]. The prevalence of dementia fast elevates from about 2-3% among those aged 70–75 years to 20–25% among those aged 85 years or more [Aarsland, 2020].

Alzheimer’s disease is the neurodegenerative origin of dementia [Chancellor et al. 2014]. Many investigations suggest that modifications in multiple genes and signaling pathways are associated in controlling the advancement of dementia. However, an inadequacy of investigation on the precise molecular mechanisms of dementia advancement limits the treatment efficacy of the disease at present. Therefore, understanding the molecular mechanisms of dementia occurrence and progression is of utmost essential for diagnosis, prognosis and targeted therapy in the future.

In recent decades, more and more researchers have devoted themselves to exploring the molecular mechanisms by which signaling pathways regulates the progression of dementia. Signaling pathways includes TGF-β signaling pathway [Kandasamy et al. 2020], TLR4/MyD88/NF-κ β signaling pathway [Wang et al. 2020], Akt/mTOR signaling pathway [Xu et al. 2019], SIRT1/PGC-1α signaling pathway [Yao et al. 2019] and VEGF–VEGFR2 signaling pathway [Wang et al. 2014] are involved in the initiation and progression of dementia. Further, a series of genes such as UBQLN2 [Deng et al. 2011], PRKAR1B [Cohn-Hokke et al. 2014], apolipoprotein E (APOE) [Regland et al. 1999], TREM2 [Guerreiro et al. 2013] and PON1 [Alam et al. 2014], have been predicted as target genes for diagnosing and treatment of dementia. However, the interactions between genes, signal pathway, miRNAs and transcription factors (TFs) warrant further comprehensive analysis.

RNA sequencing has increasingly become a promising tool in studying neuropsychiatric disorders [Hardwick et al. 2019]. These genomic data offer possibilities for identifying certain disease related genes. Recently, many investigations have been performed according to expression profiling by high throughput sequencing data profiles to elucidate the pathogenesis of dementia [Rajkumar et al. 2020].

In the current study, we downloaded expression profiling by high throughput sequencing dataset, GSE153960 [Prudencio et al. 2020], from the NCBI Gene Expression Omnibus database (NCBI GEO) (http://www.ncbi.nlm.nih.gov/geo/) [Clough and Barrett, 2016]), including gene expression data from 263 patients with dementia and 277 healthy controls. We identified differentially expressed genes (DEGs) using the limma R bioconductor package with standard data processing. To further understand the function of genes, Gene Ontology (GO) enrichment analysis, REACTOME pathway enrichment analysis, protein-protein interaction (PPI) network construction and analysis, module analysis, miRNA-hub regulatory gene network construction and analysis and TF-hub regulatory gene network construction and analysis were performed in sequence. Finally, selected hub genes were verified by receiver operating characteristic curve (ROC) analysis. We anticipated that these investigations will implement further insight of dementia pathogenesis and development at the molecular level.

## Material and Methods

### RNA sequencing data

The expression profiling by high throughput sequencing dataset (GEO access number: GSE153960, contributed by [Prudencio et al. 2020], were downloaded from the NCBI GEO, which was currently the largest fully public gene expression resource. There were a total of 540 samples in this analysis, including 263 patients with dementia and 277 healthy controls.

### Identification of DEGs

The DEGs were identified by using limma package in R between patients with dementia and healthy controls [Ritchie et al. 2015], with the criteria of |log fold change (FC)| > 0.6 for up regulated genes, |log fold change (FC)| < -0.7 for down regulated genes and p-value < 0.05 followed by multiple-testing correction using the Benjamini-Hochberg procedure to obtain the adjusted p-value [Hardcastle, 2016].

### GO and REACTOME pathway enrichment analysis

ToppGene (ToppFun) (https://toppgene.cchmc.org/enrichment.jsp) [Chen et al 2009] is a tool which provides a comprehensive set of functional annotation tools for researchers to investigate the biological meaning of genes. Identified DEGs were investigated further using ToppGene, Gene Ontology (GO) (http://geneontology.org/) [Thomas, 2017] and REACTOME (https://reactome.org/) [Fabregat et al. 2018] pathway enrichment analyses. P<0.05 and gene counts of >5 were considered to indicate a statistically significant difference in the functional enrichment analysis.

### PPI network construction and module analysis

Identified DEGs were mapped into the online Search Tool for the Retrieval of Interacting Genes (STRING) database [Szklarczyk et al. 2011] to evaluate the interactive relationships among the DEGs. Interactions with a combined score >0.4 were defined as statistically significant. Cytoscape software (version 3.8.2) (www.cytoscape.org) [Shannon et al. 2003] was used to visualize the integrated regulatory networks. According to the node degree [Przulj et al 2004], betweenness centrality [Nguyen et al 2011], stress centrality [Shi and Zhang, 2011] and closeness centrality [Fadhal et al 2014] in the Cytoscape plugin Network Analyzer, the top five ranked genes were defined as hub genes. . The Cytoscape plugin PEWCC1 (http://apps.cytoscape.org/apps/PEWCC1) [Zaki et al 2013] was used to further detect deeper connected regions within the PPI network.

### MiRNA-hub gene regulatory network construction

MiRNA-hub gene regulatory was constructed employing the hub genes. MiRNA targets were extracted from miRNet database (https://www.mirnet.ca/) [Fan and Xia, 2018]. The regulatory interactions between miRNA and hub genes were obtained. The miRNA-target regulatory networks were visualized by Cytoscape.

### TF-hub gene regulatory network construction

TF-hub gene regulatory was constructed employing the hub genes. TF targets were extracted from NetworkAnalyst database (https://www.networkanalyst.ca/) [Zhou et al 2019. The regulatory interactions between TF and hub genes were obtained. The TF-target regulatory networks were visualized by Cytoscape.

### Validation of hub genes by receiver operating characteristic curve (ROC) analysis

The multivariate modeling with selected hub genes were used to identify biomarkers with high sensitivity and specificity for dementia diagnosis. Used one data as training set and other as validation set iteratively. The receiver operator characteristic curves were plotted and area under curve (AUC) was determined independently to check the conduct of each model using the R packages “pROC” [Robin et al 2011]. An AUC 9 marked that the model had a positive fitting effect.

## Results

### Identification of DEGs

There were 263 patients with dementia and 277 healthy controls in our present study. Via Limma in R bioconductor package, we identified 948 DEGs from GSE153960. Among them, 475 genes were up regulated; while 473 were down regulated (Table 1). A heatmap of the up regulated and down regulated genes, was produced using the gplots package and R software. A volcano plot of all of the DEGs was generated using ggplot2 package and R software. The heatmap and volcano plot are shown in Fig. 1 and and Fig. 2, respectively.

**Fig. 1.**
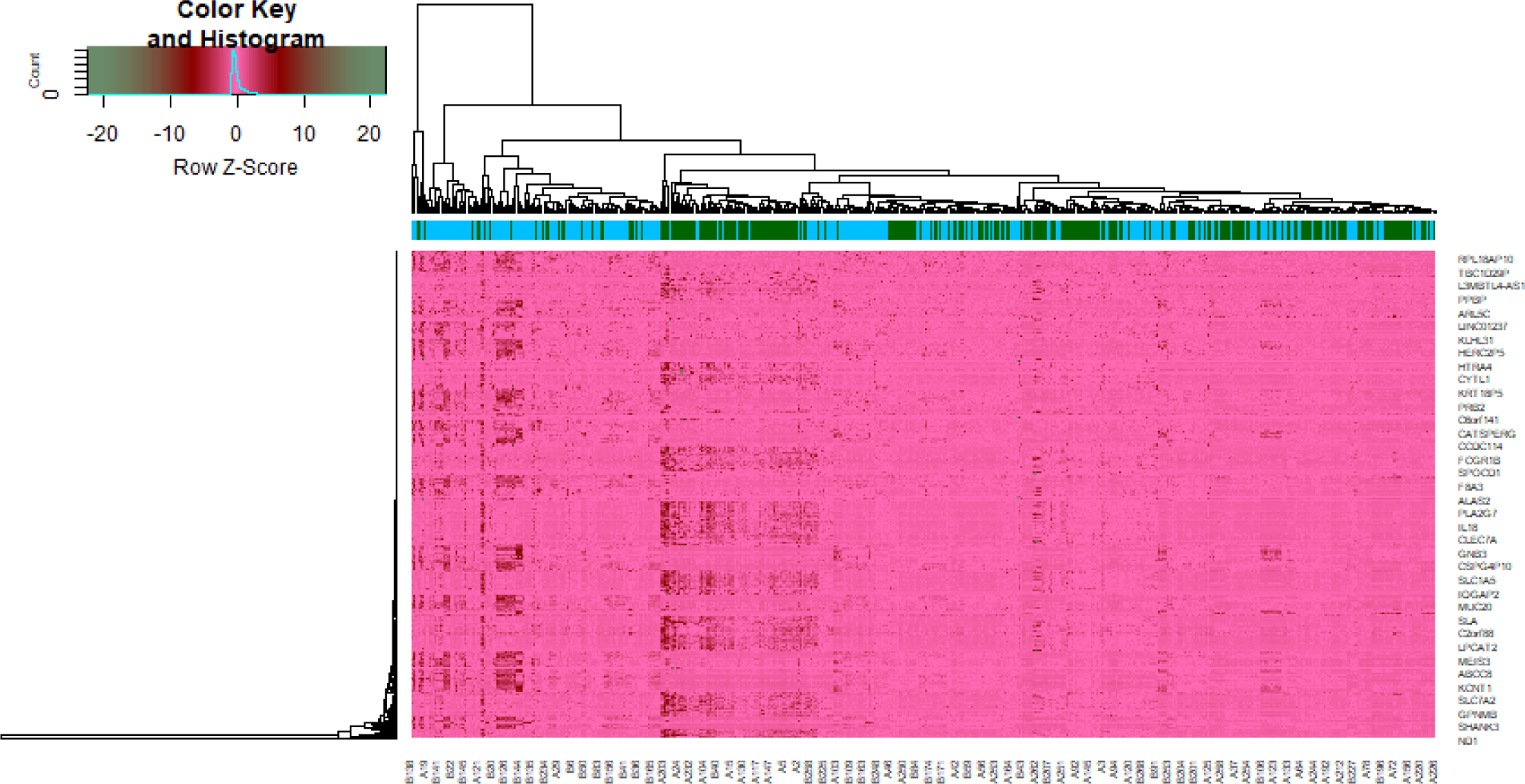
Heat map of differentially expressed genes. Legend on the top left indicate log fold change of genes. (A1 – A277= healthy controls; B1 – B263 = patients with dementia)

**Fig. 2.**
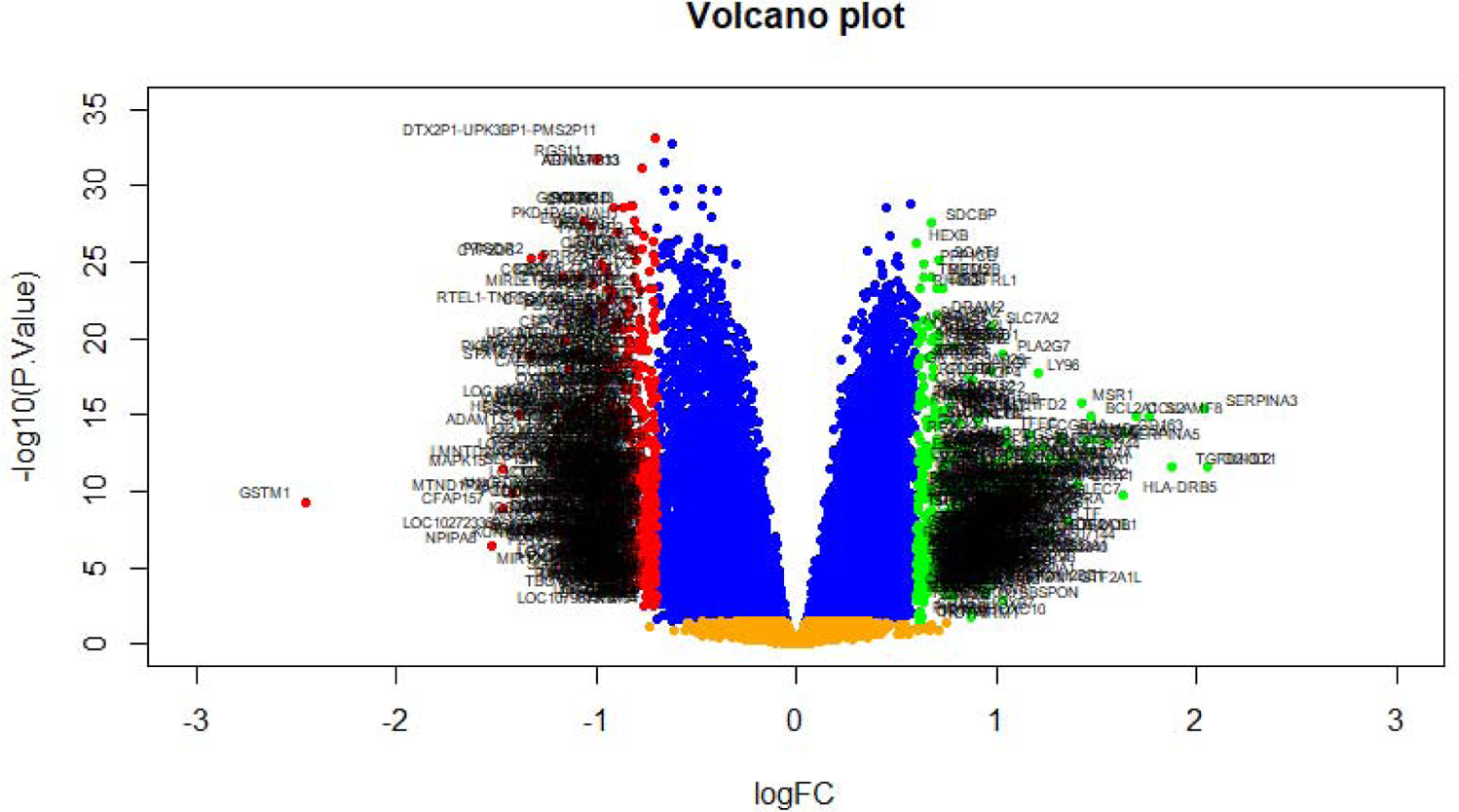
Volcano plot of differentially expressed genes. Genes with a significant change of more than two-fold were selected. Green dot represented up regulated significant genes and red dot represented down regulated significant genes.

**Table 1.**
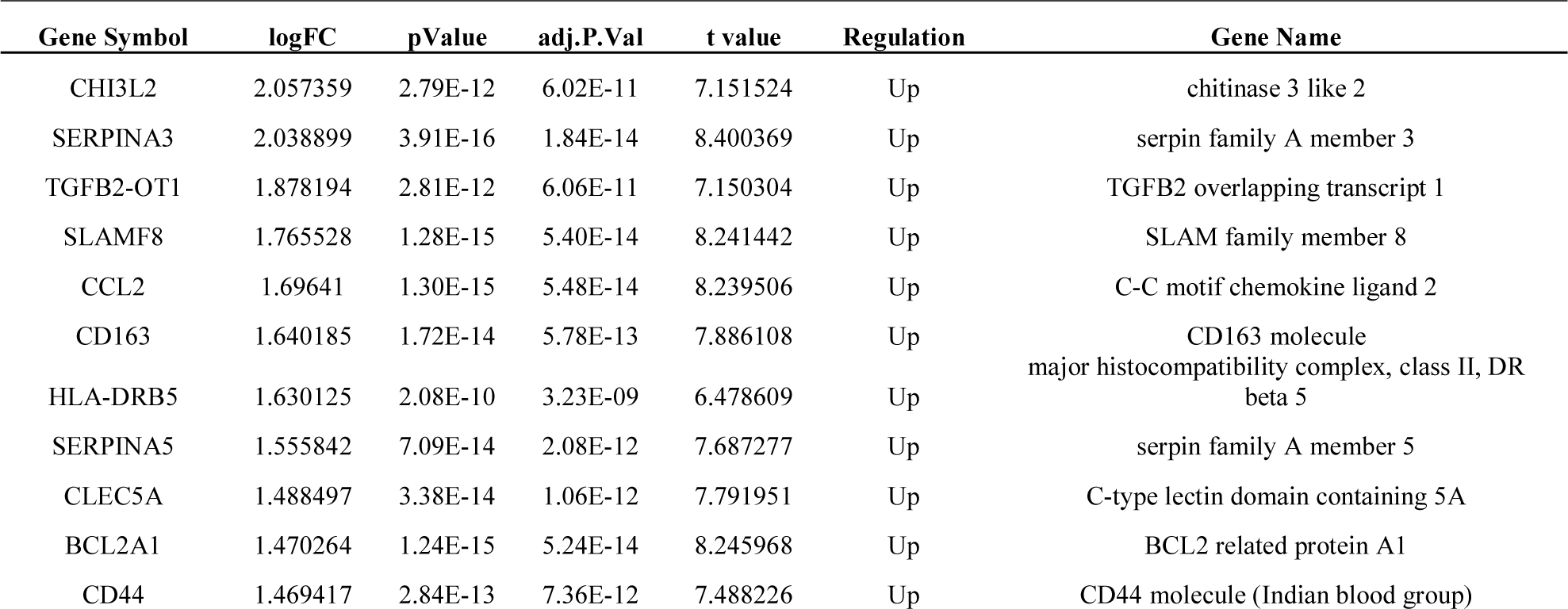
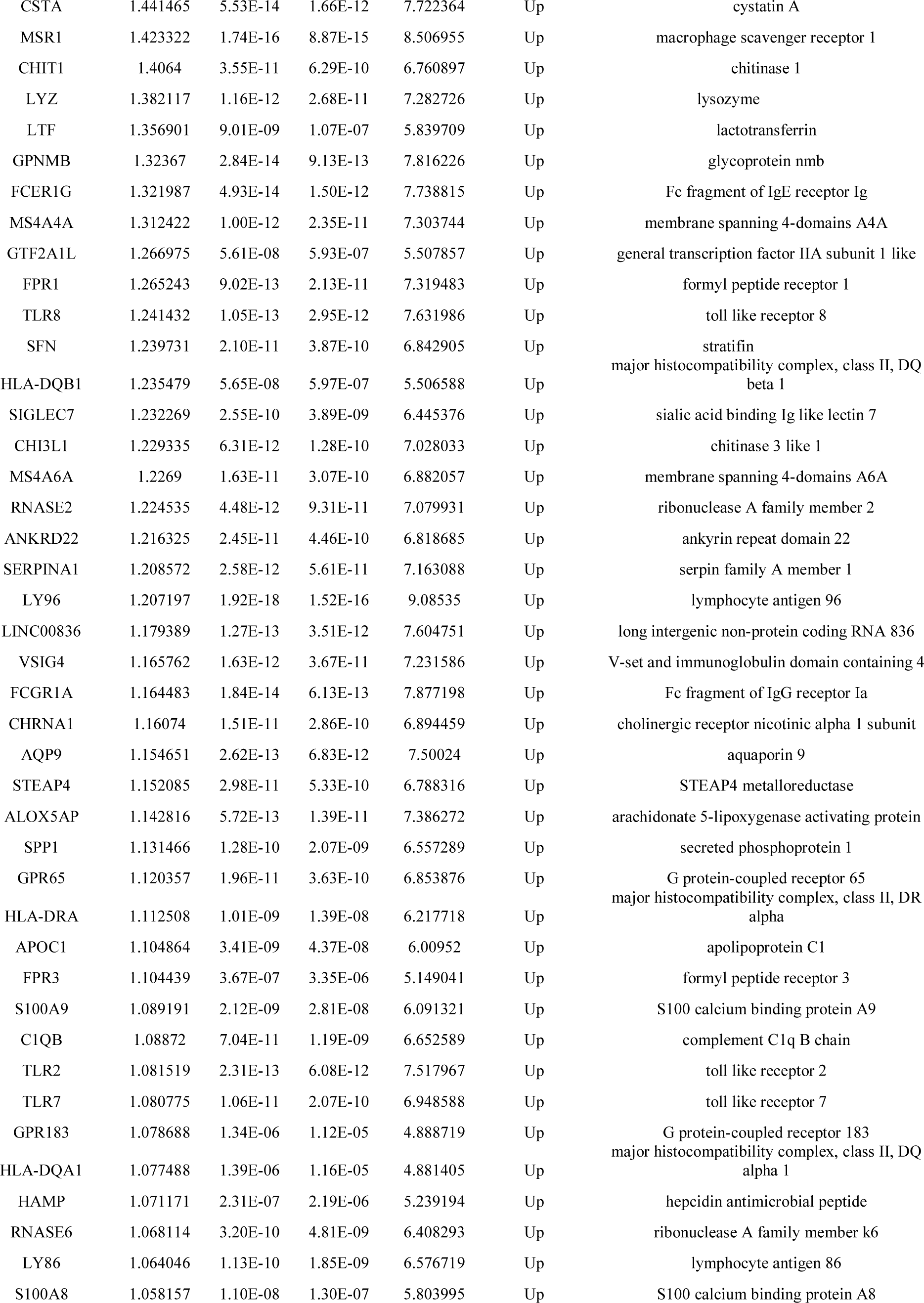
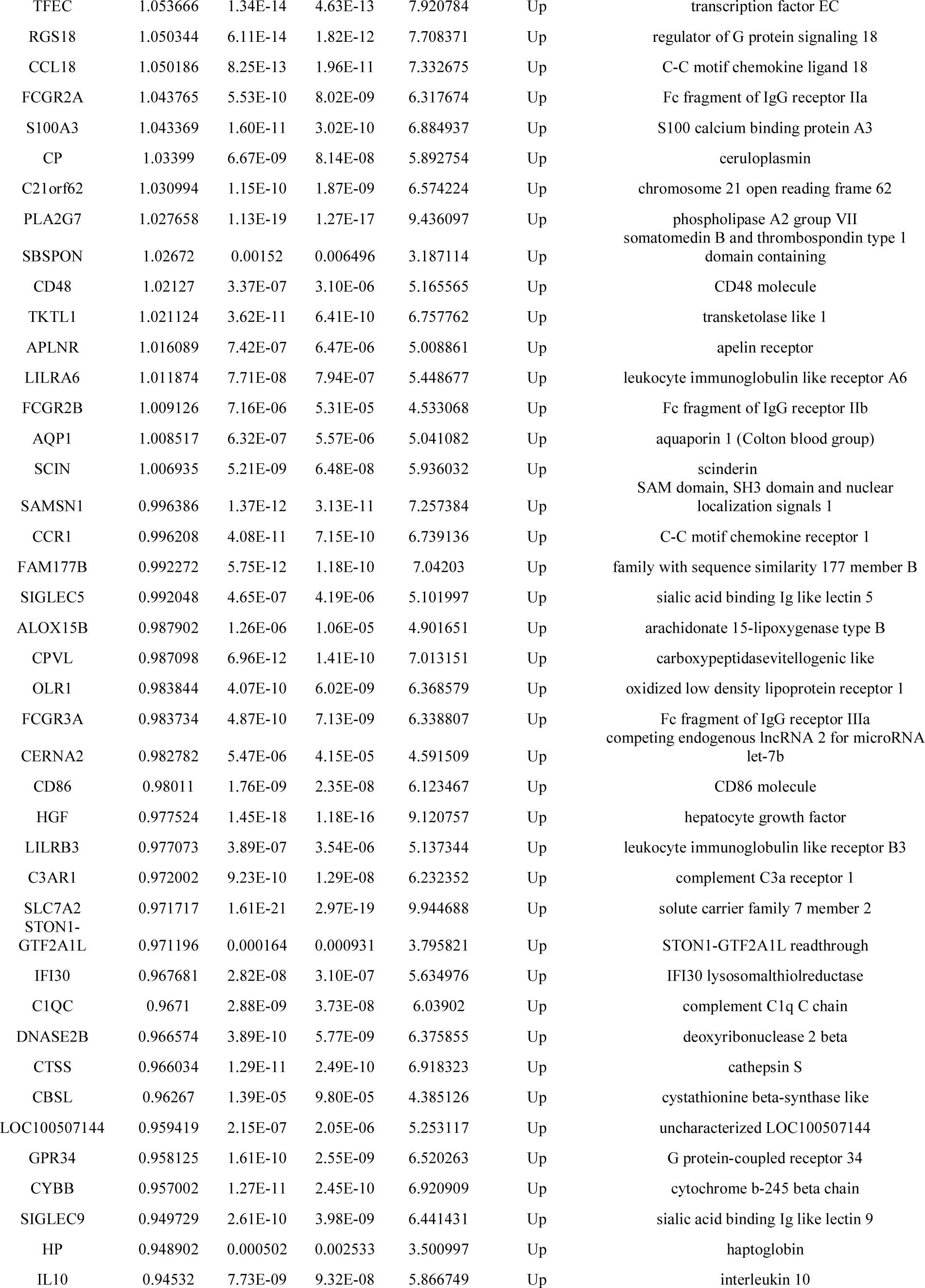
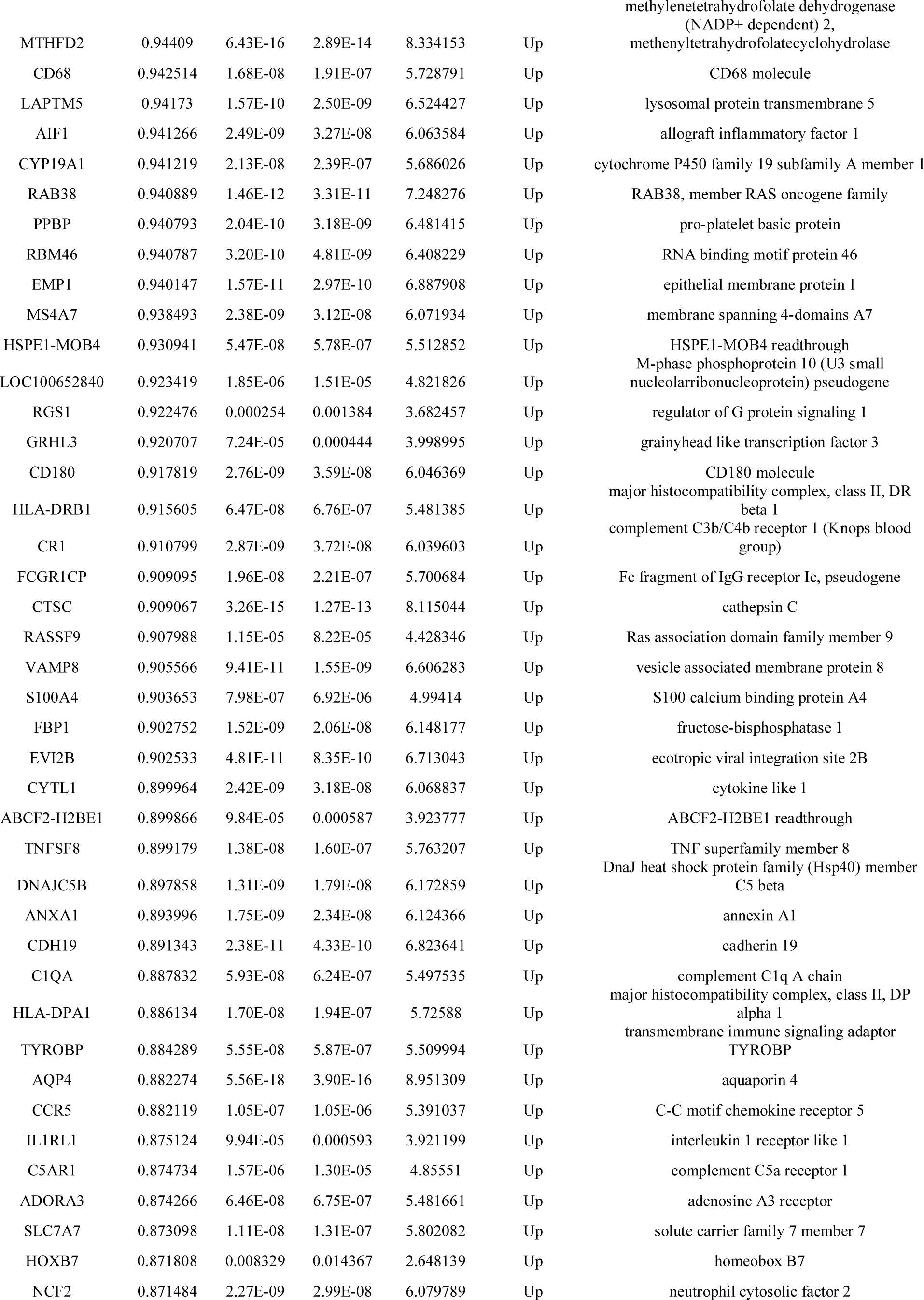
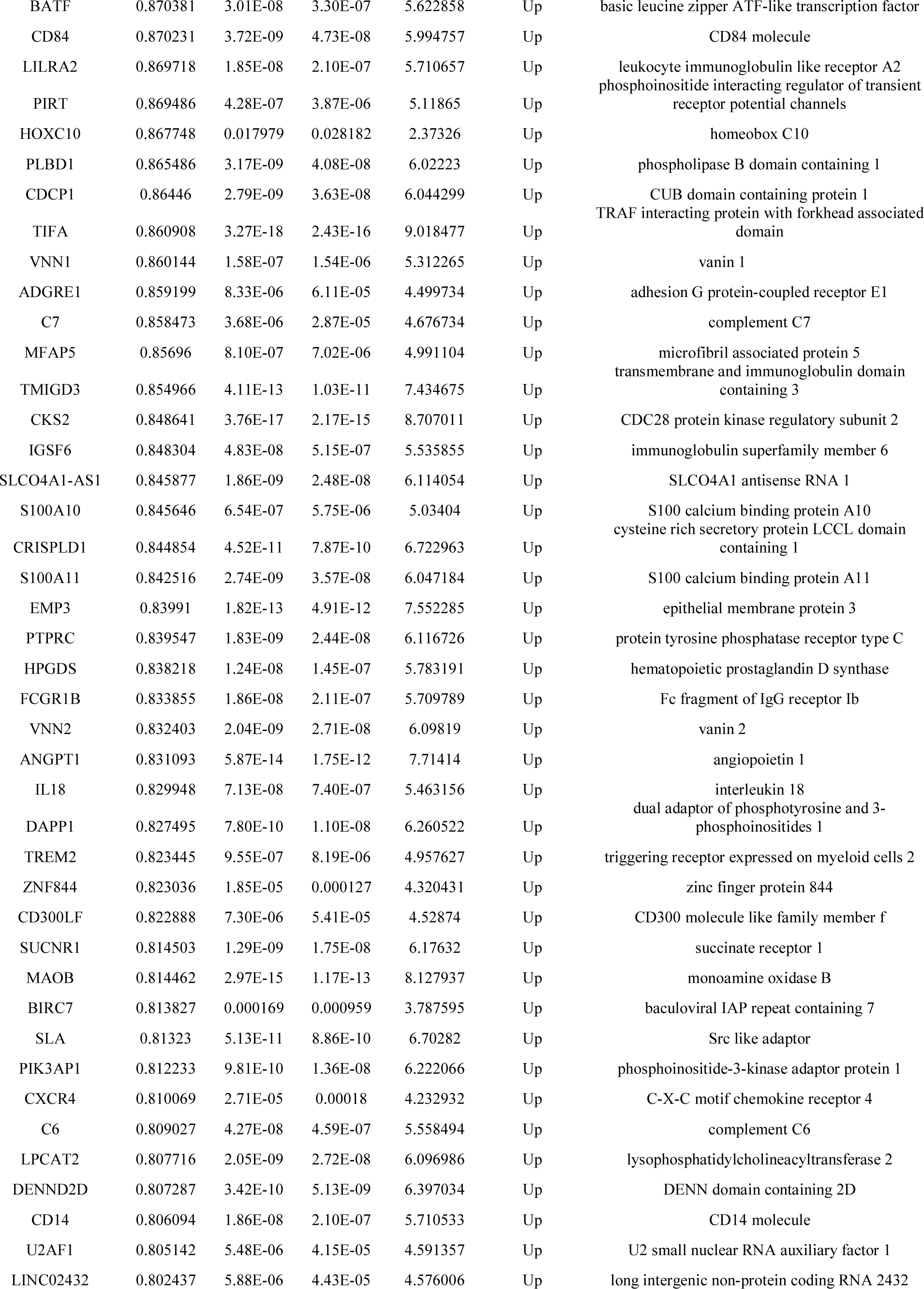
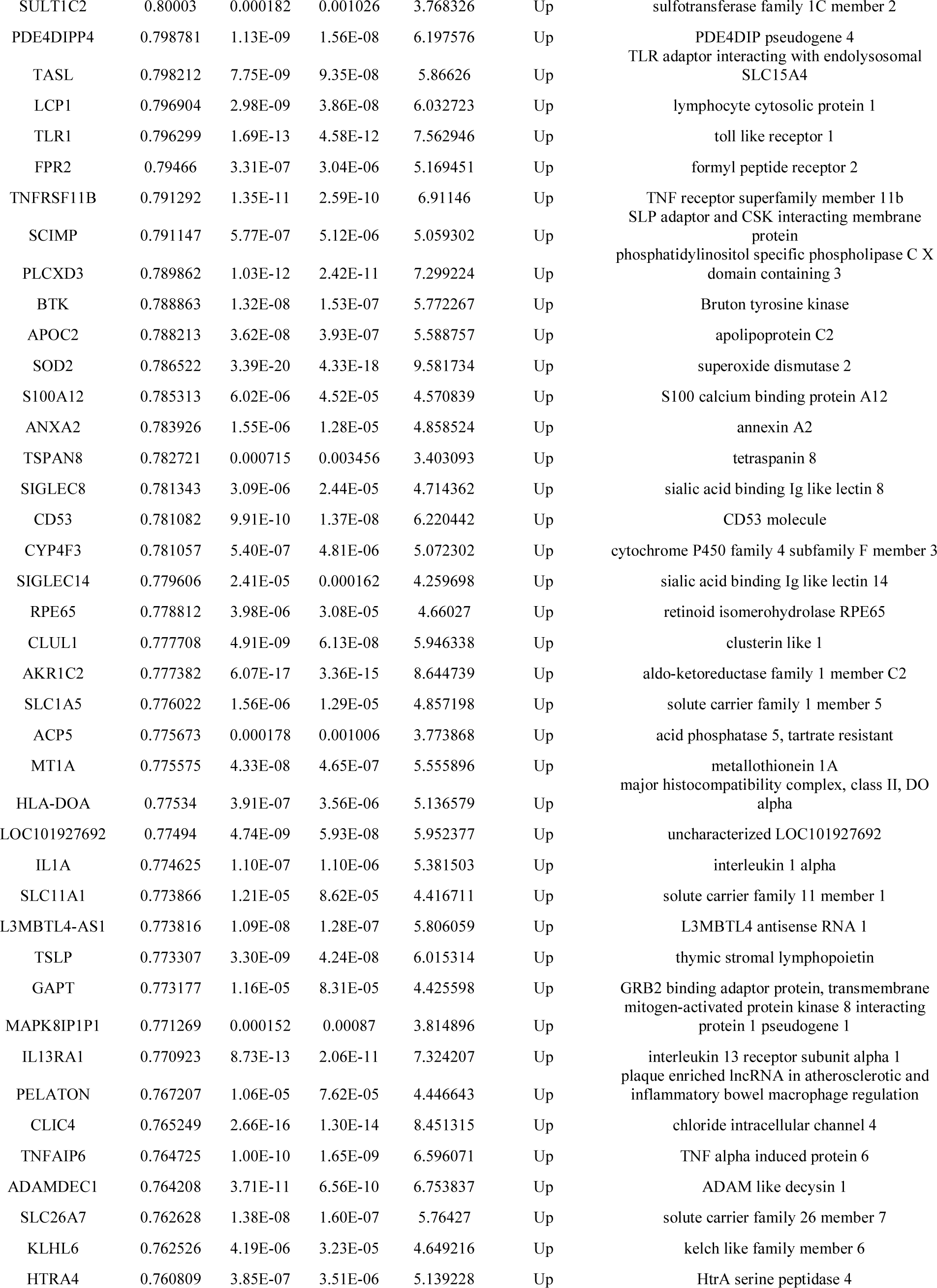
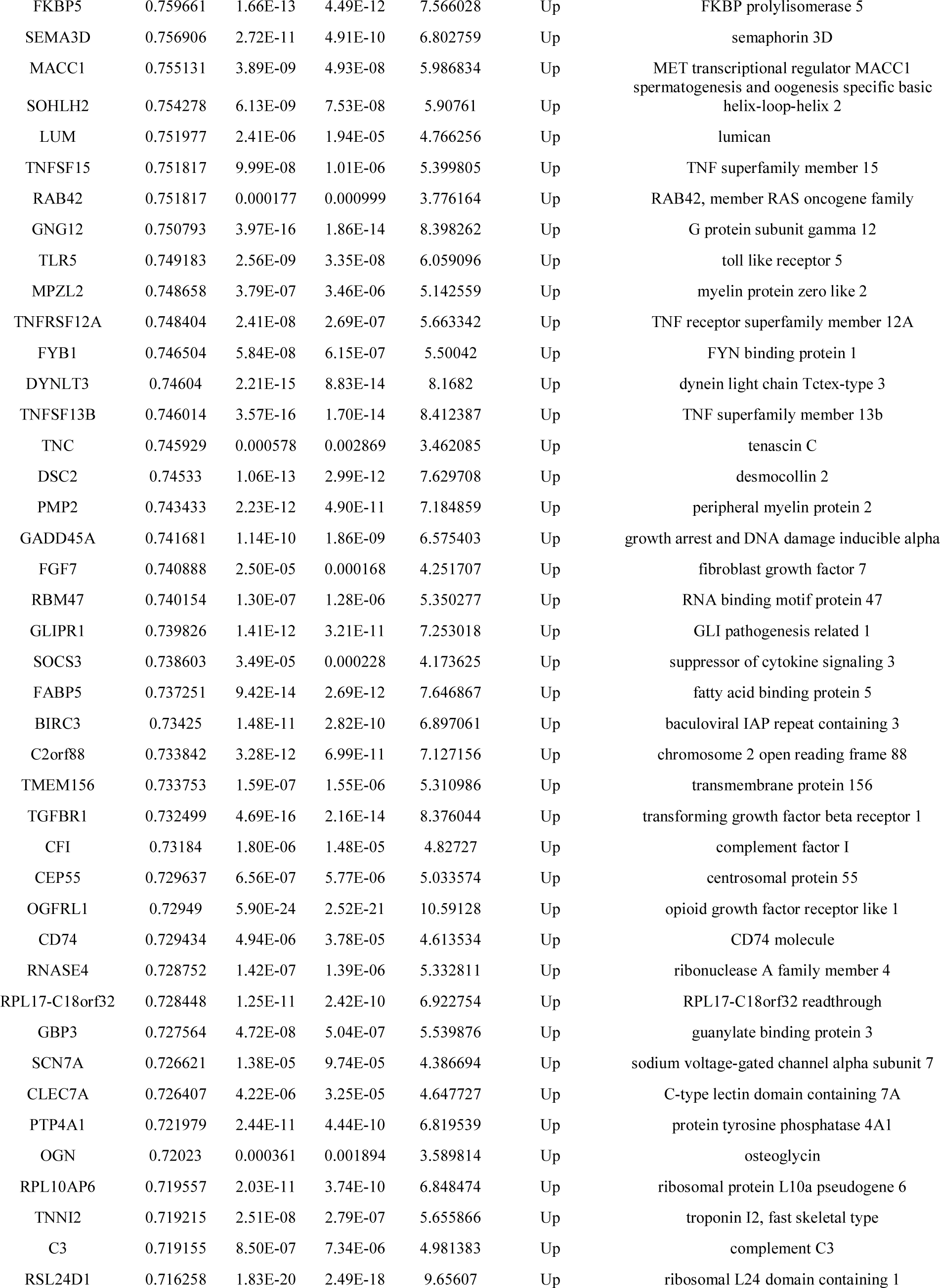
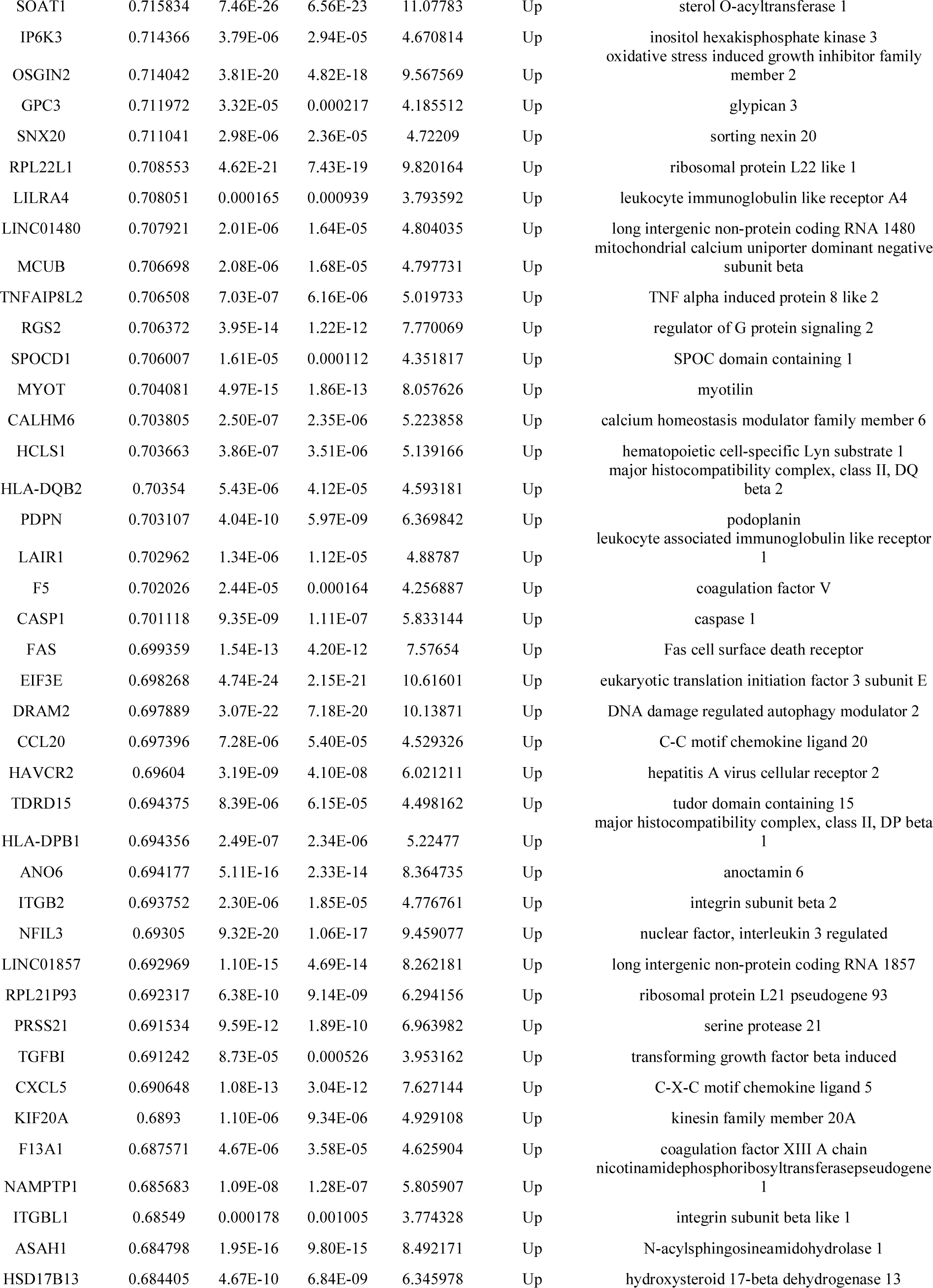
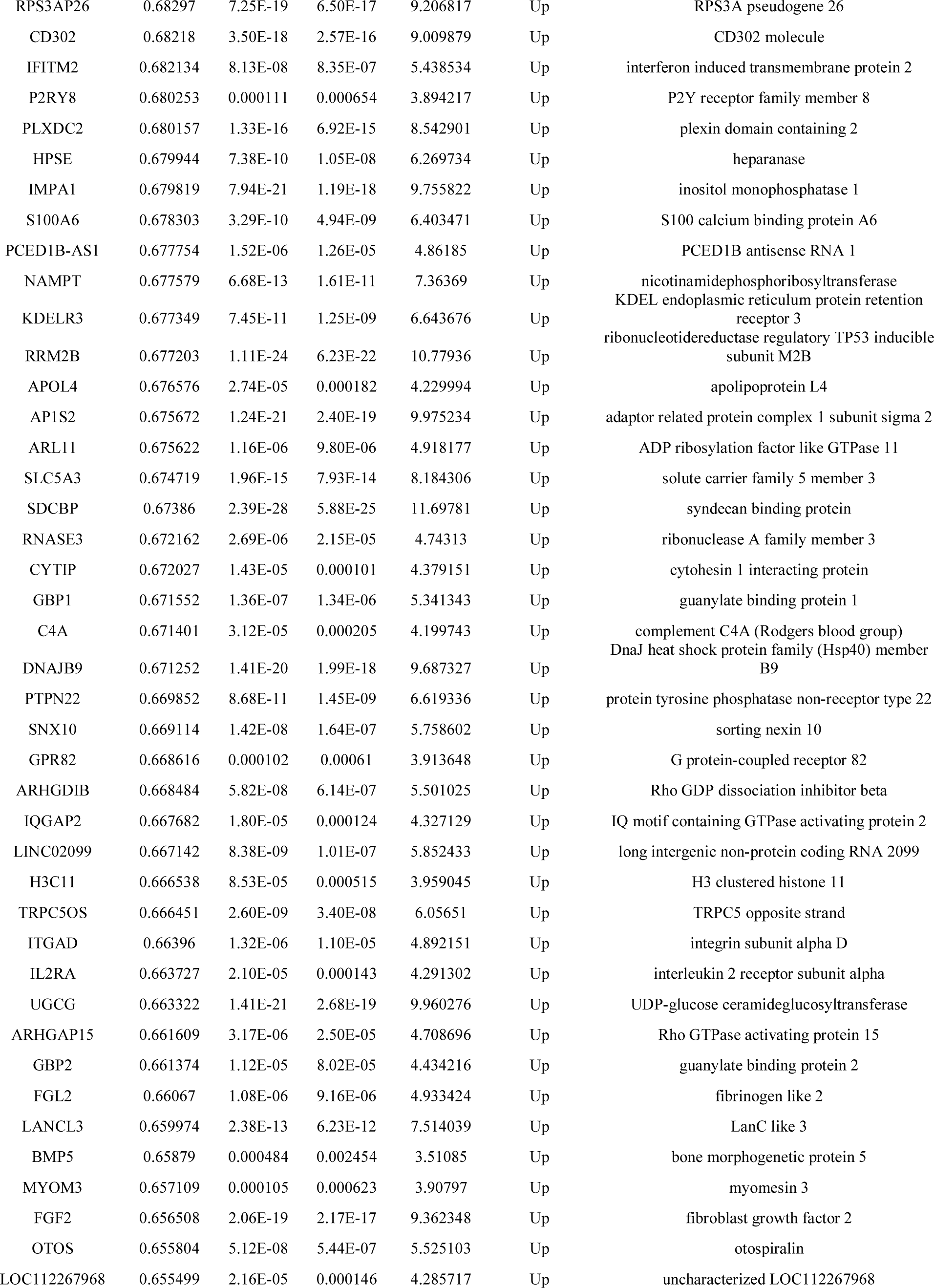
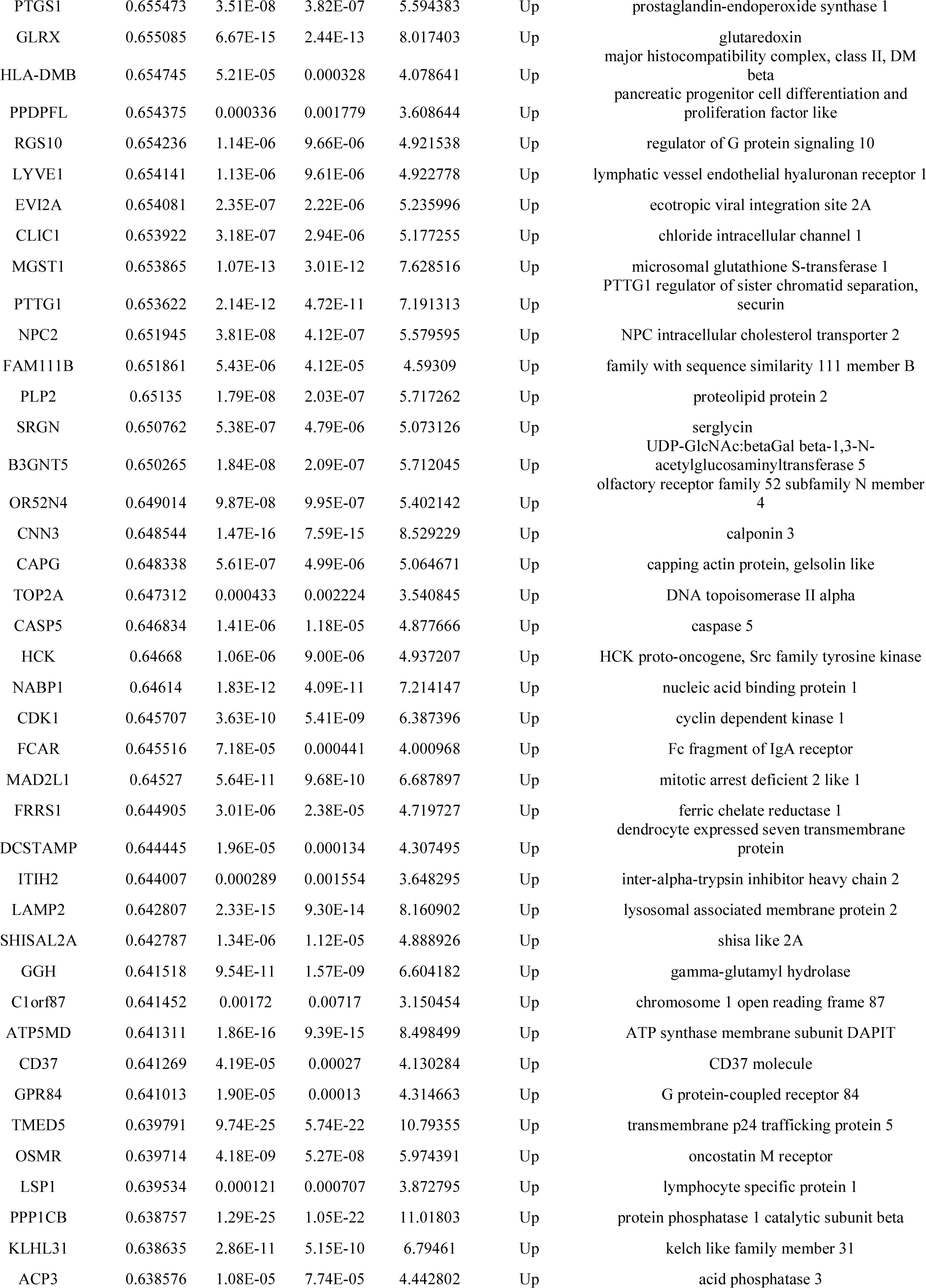
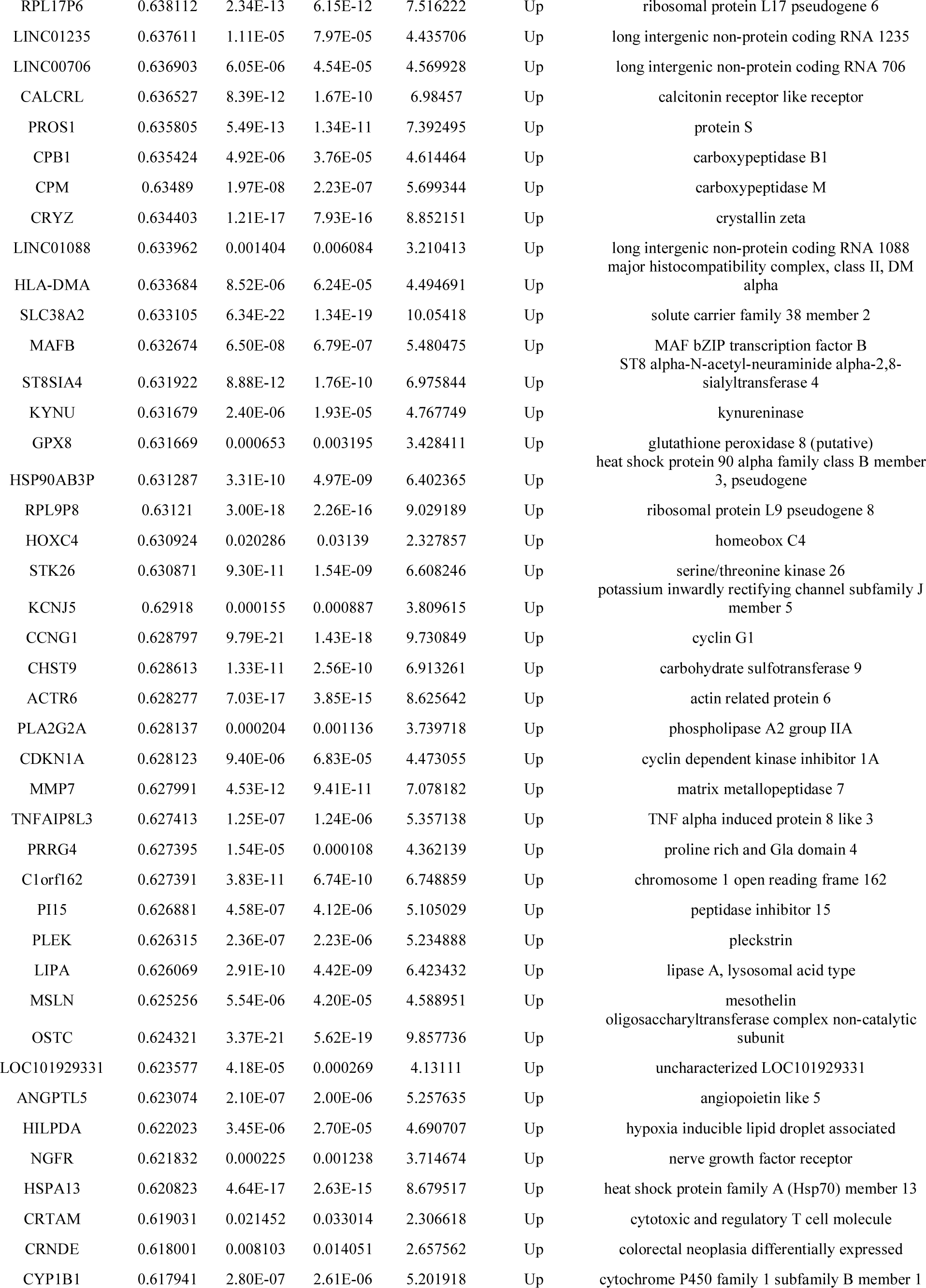
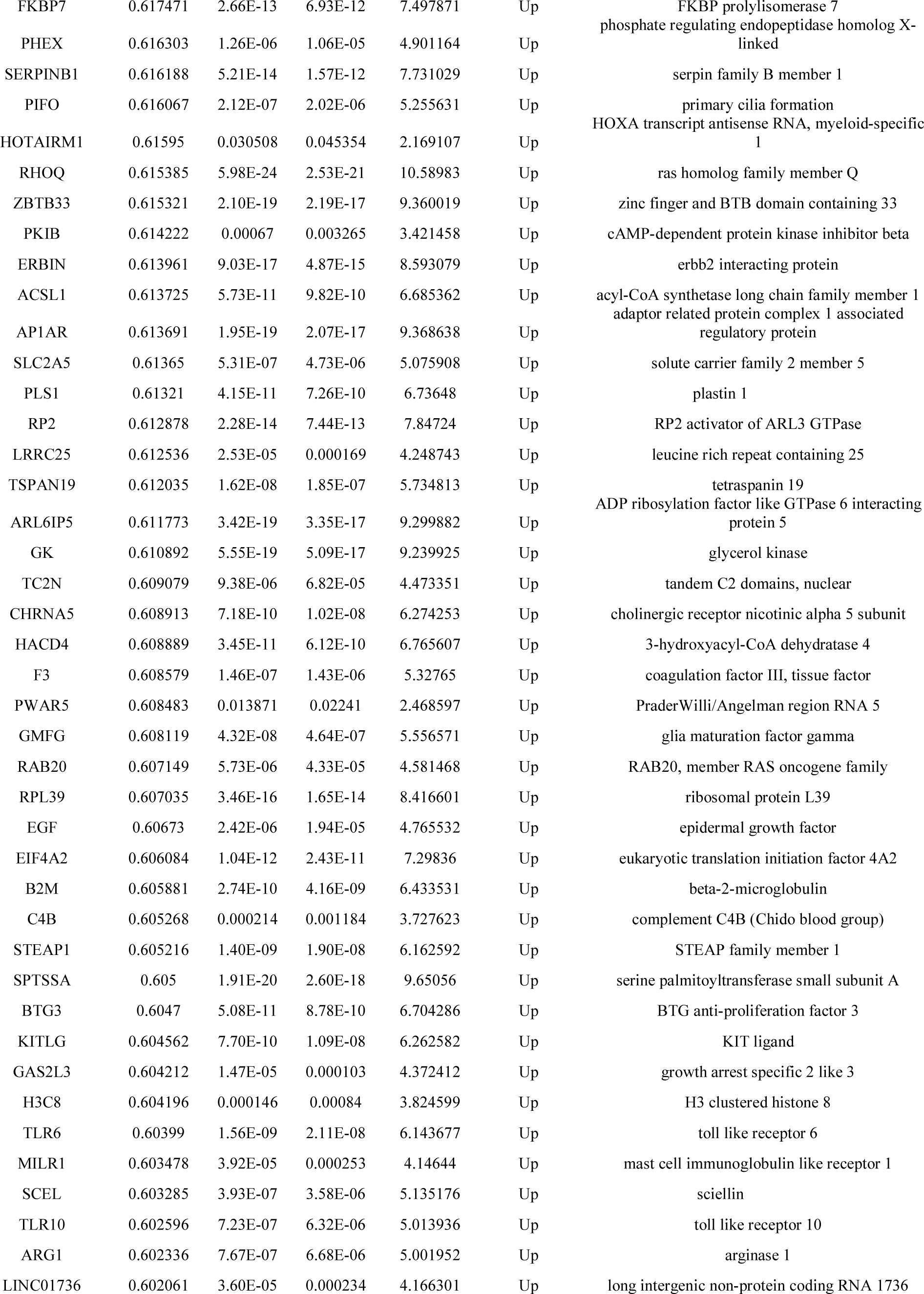
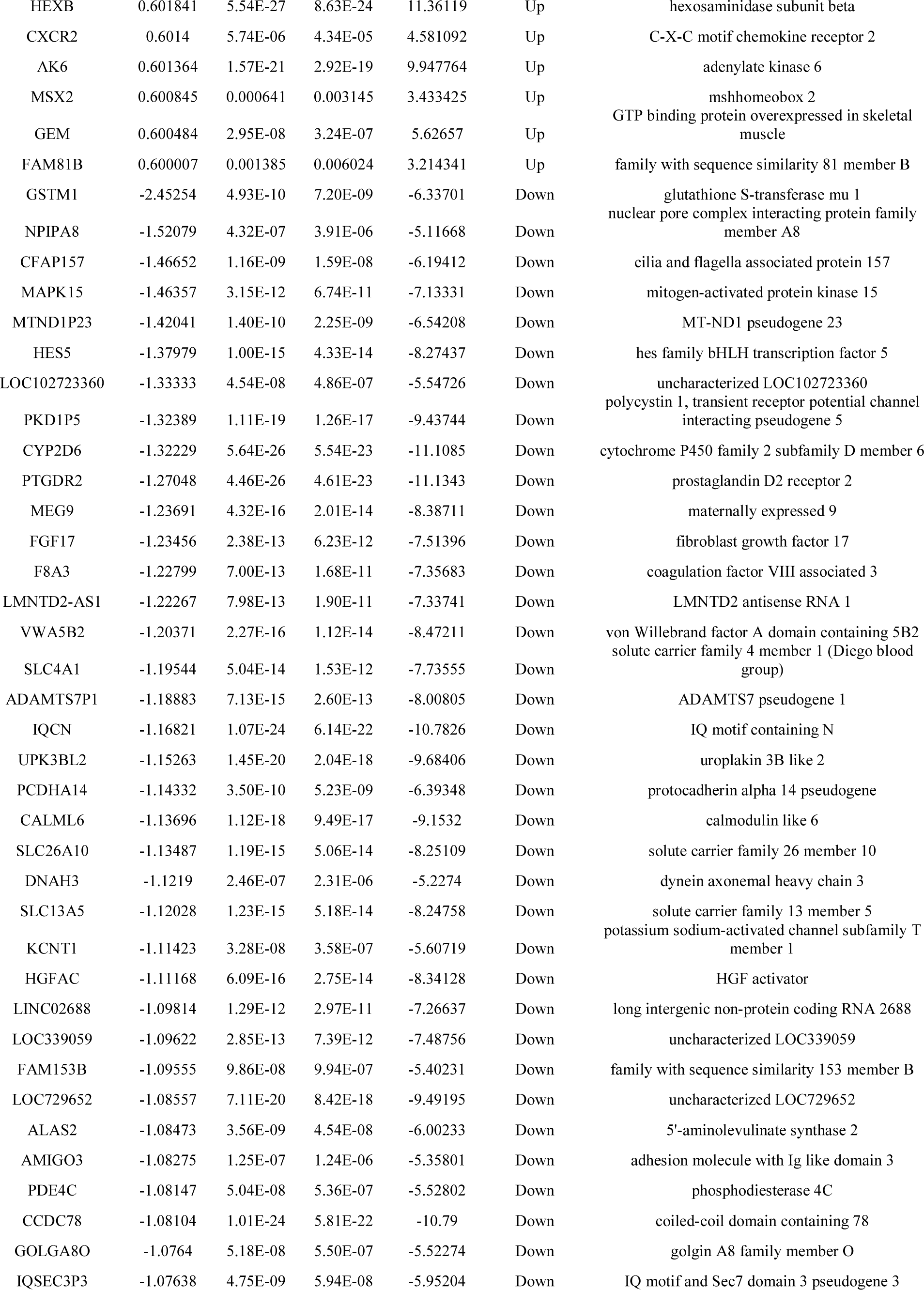
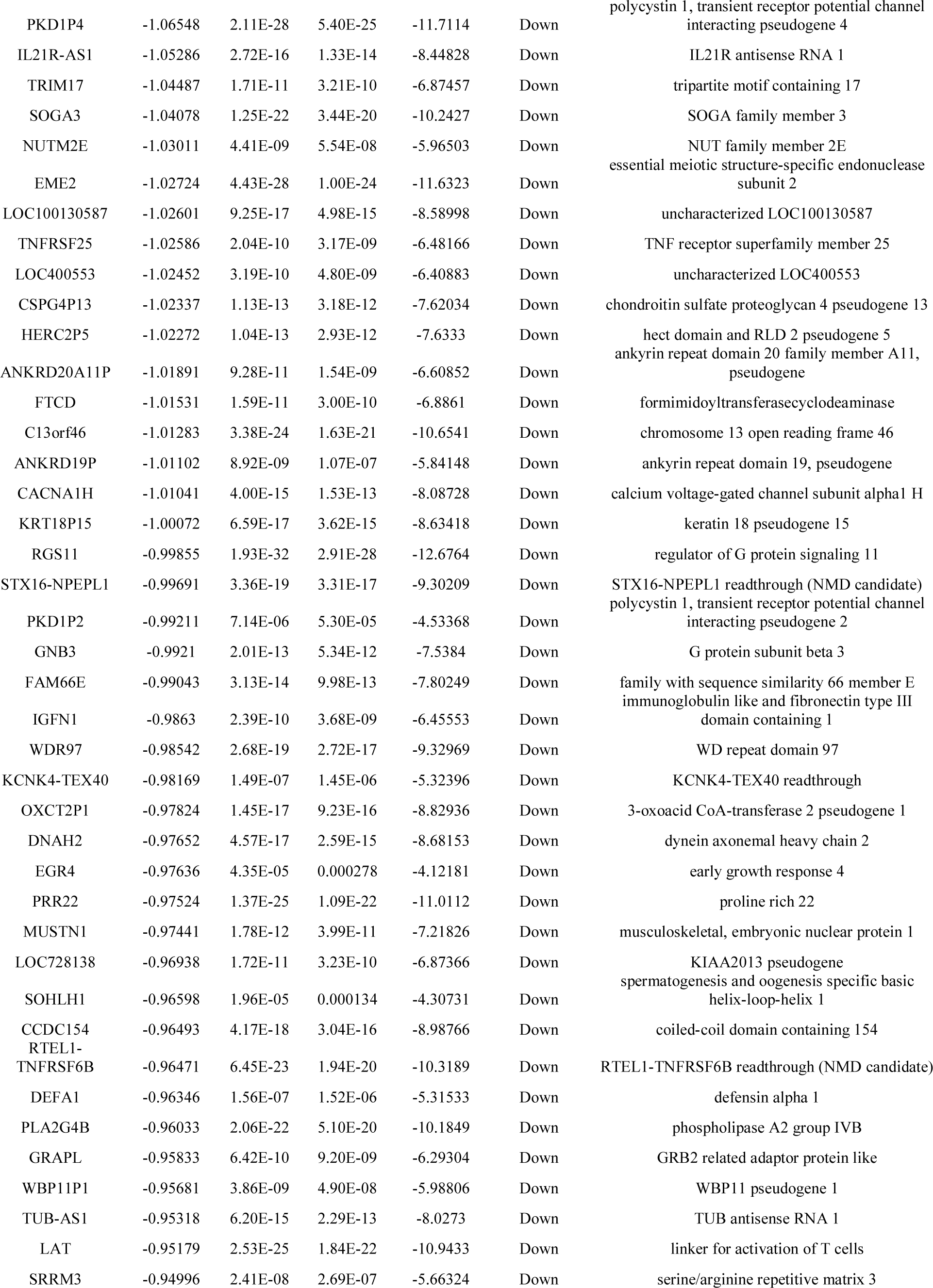
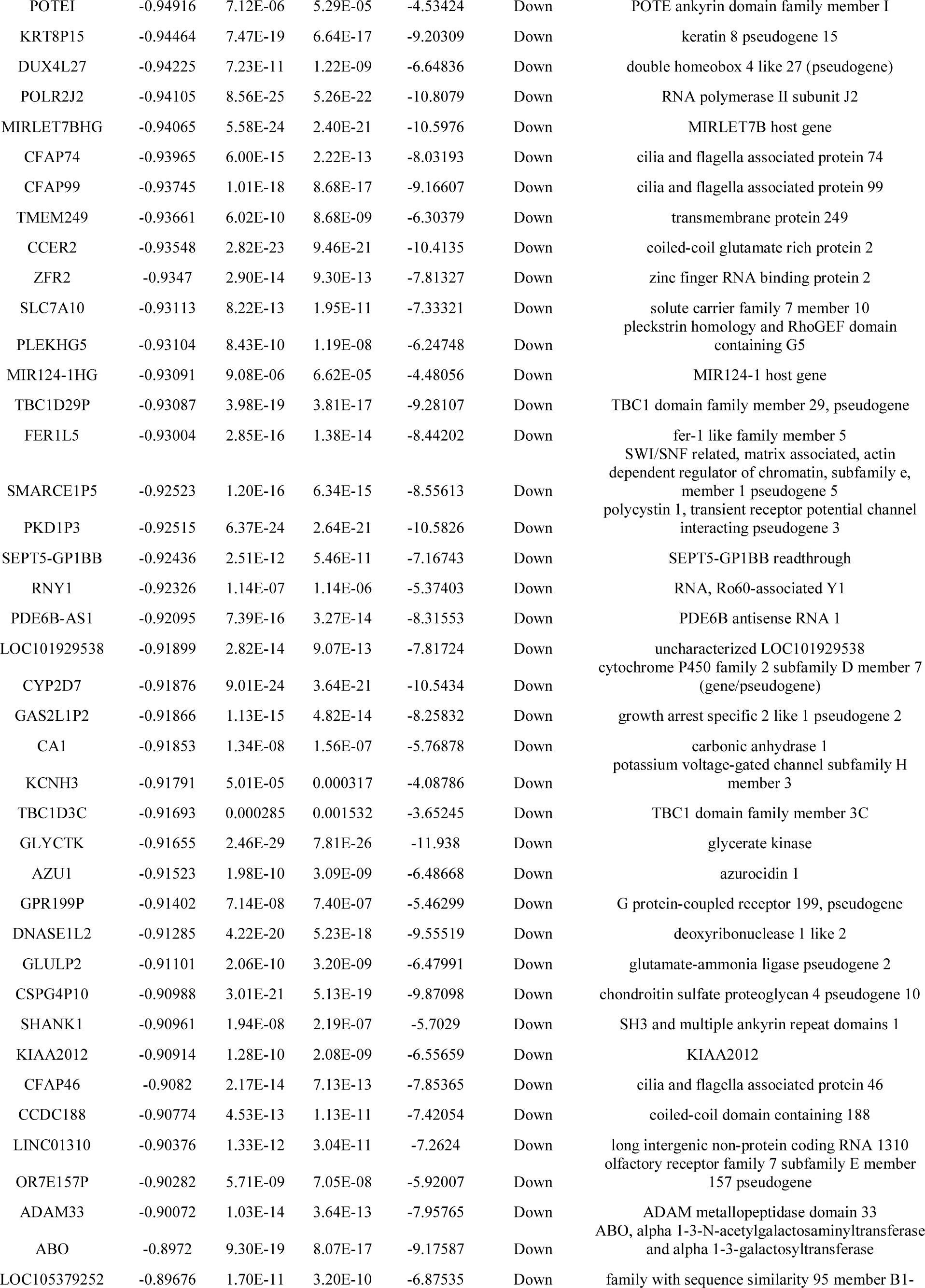
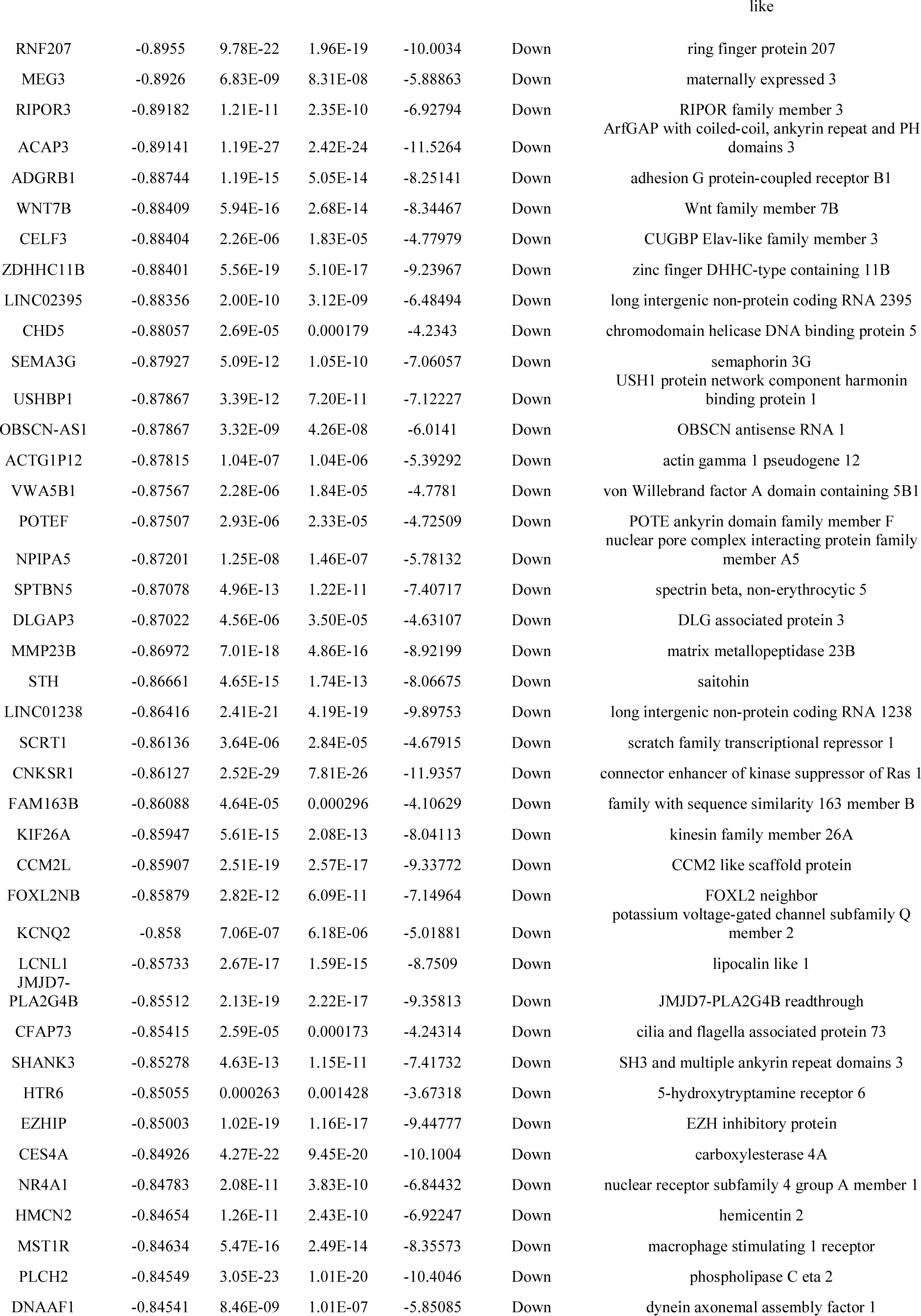
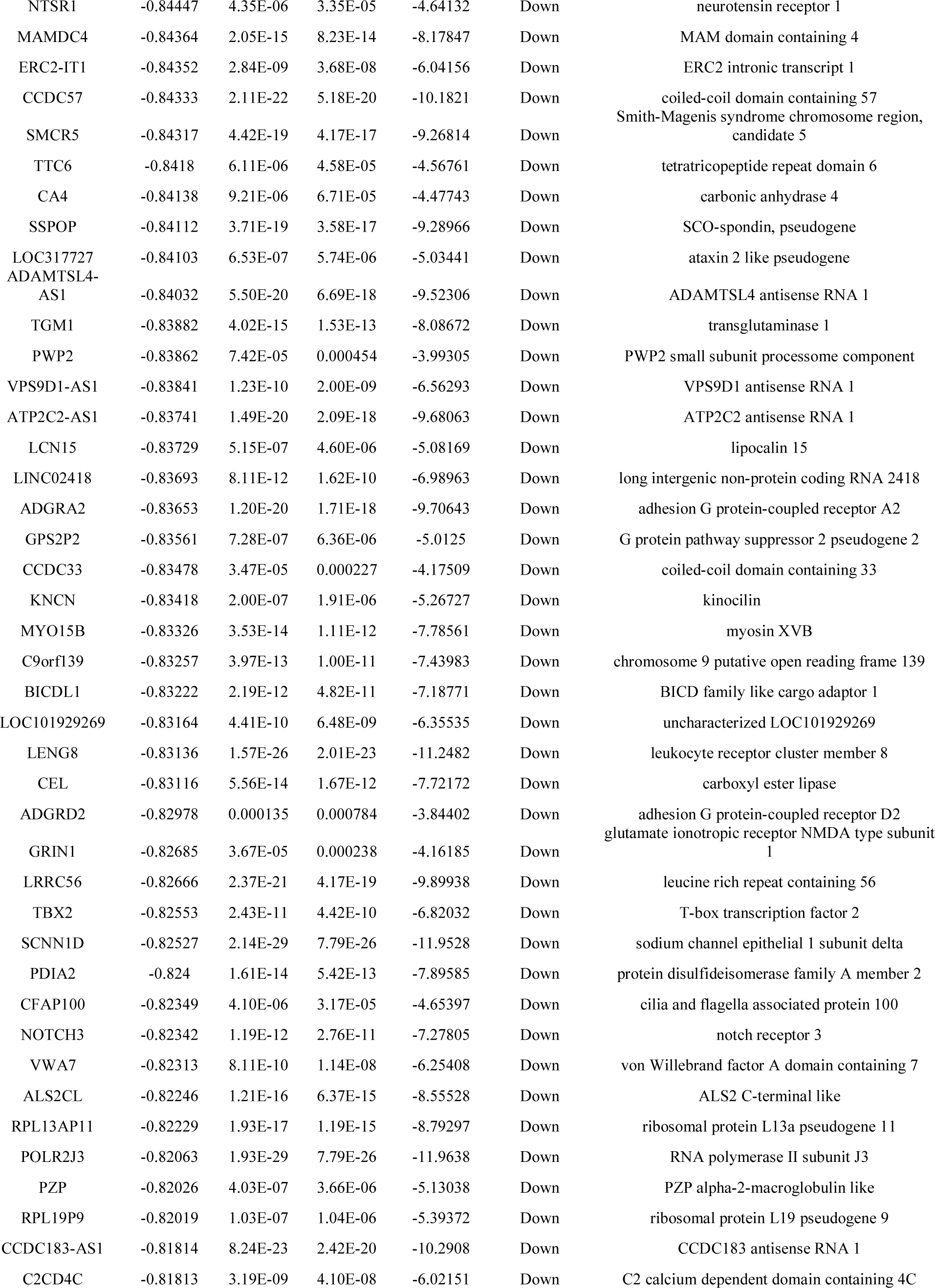
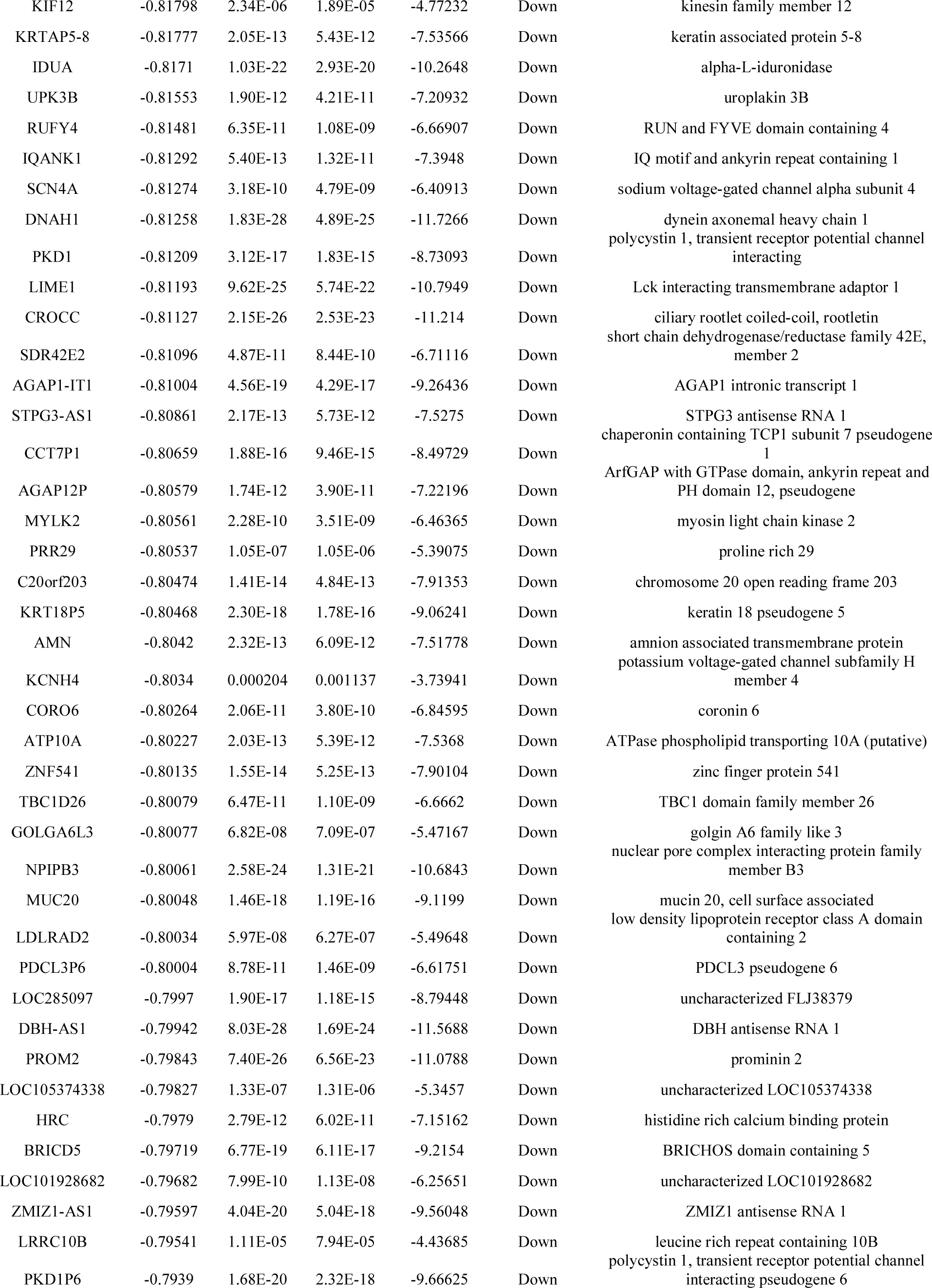
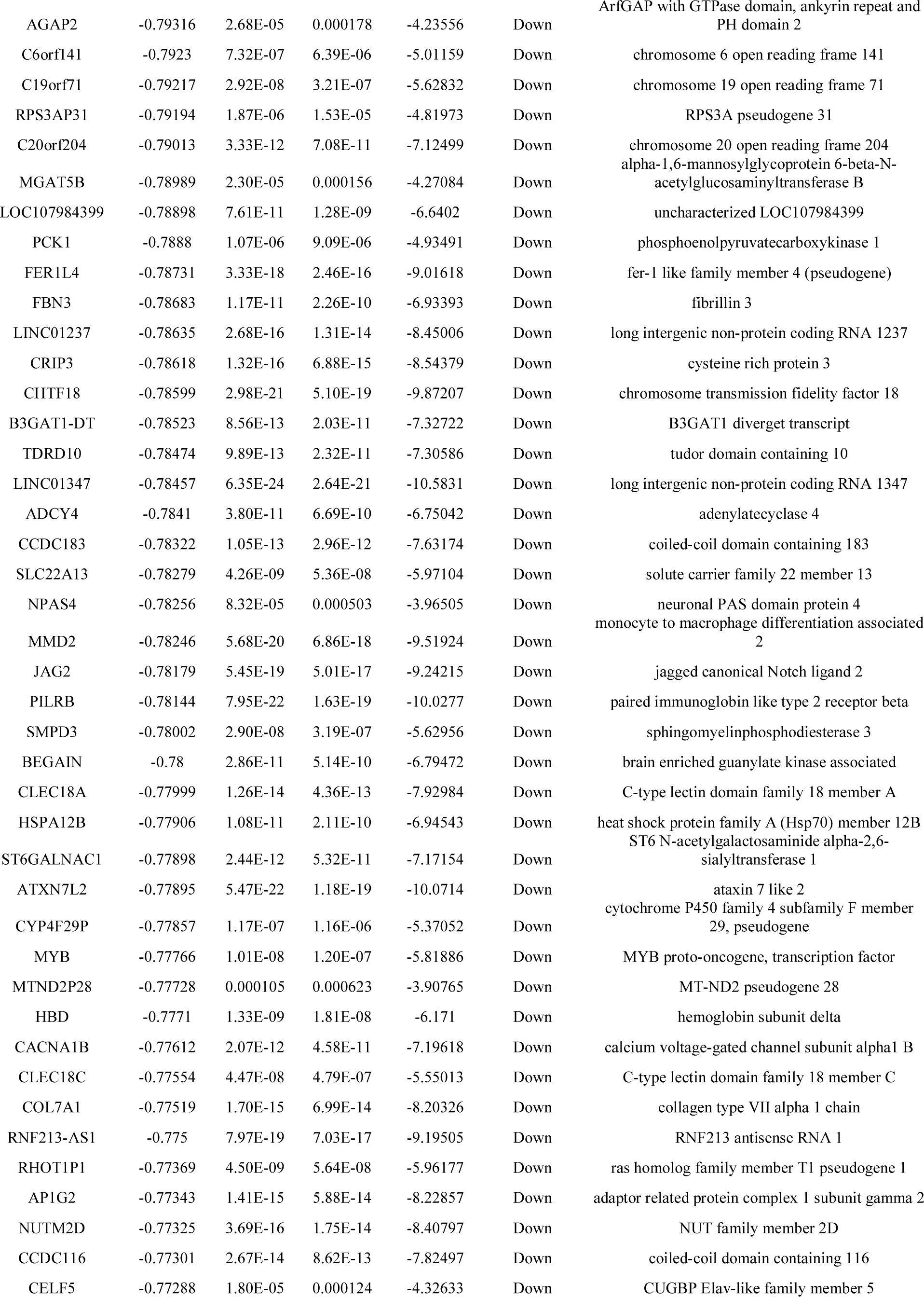
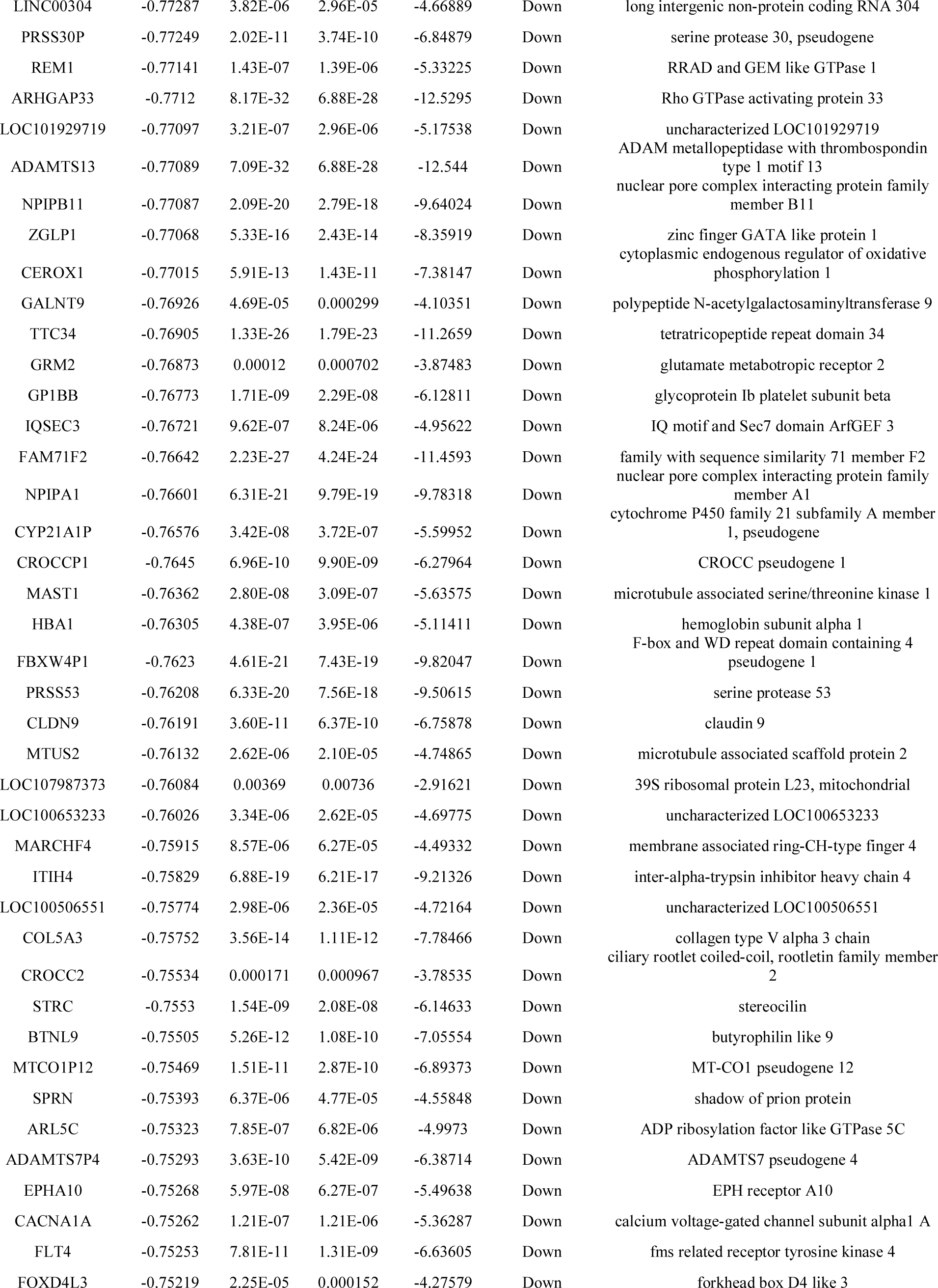
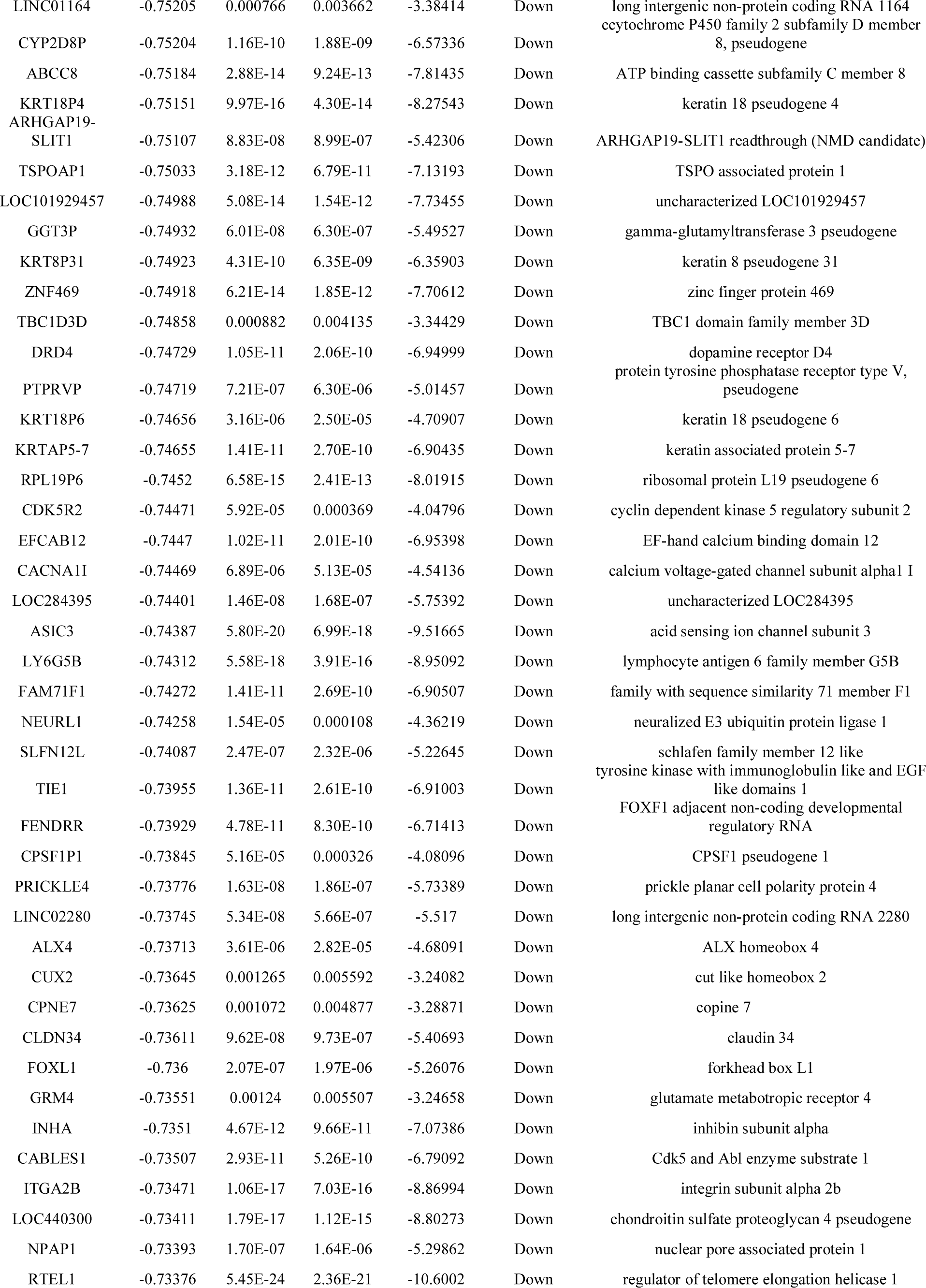
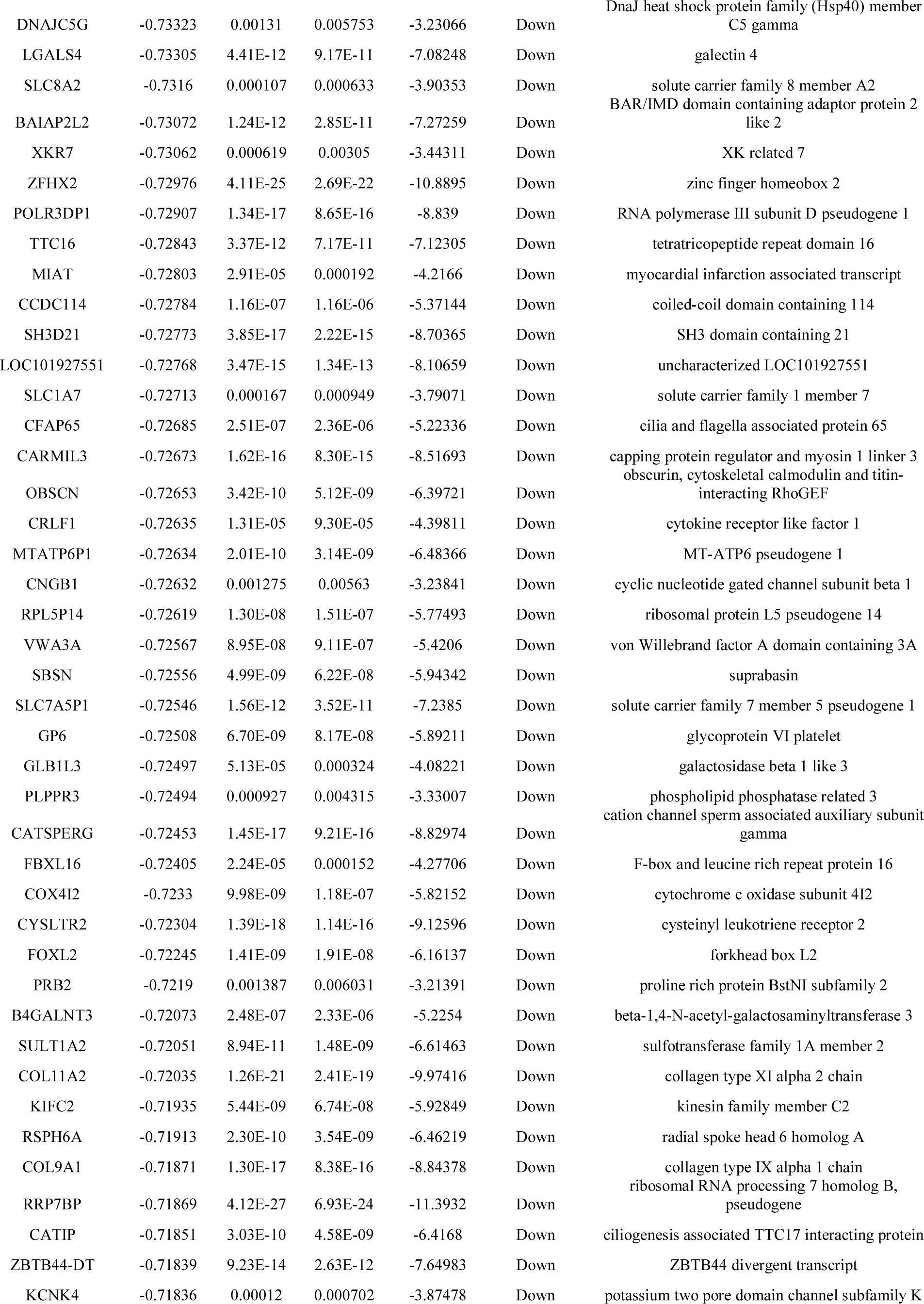
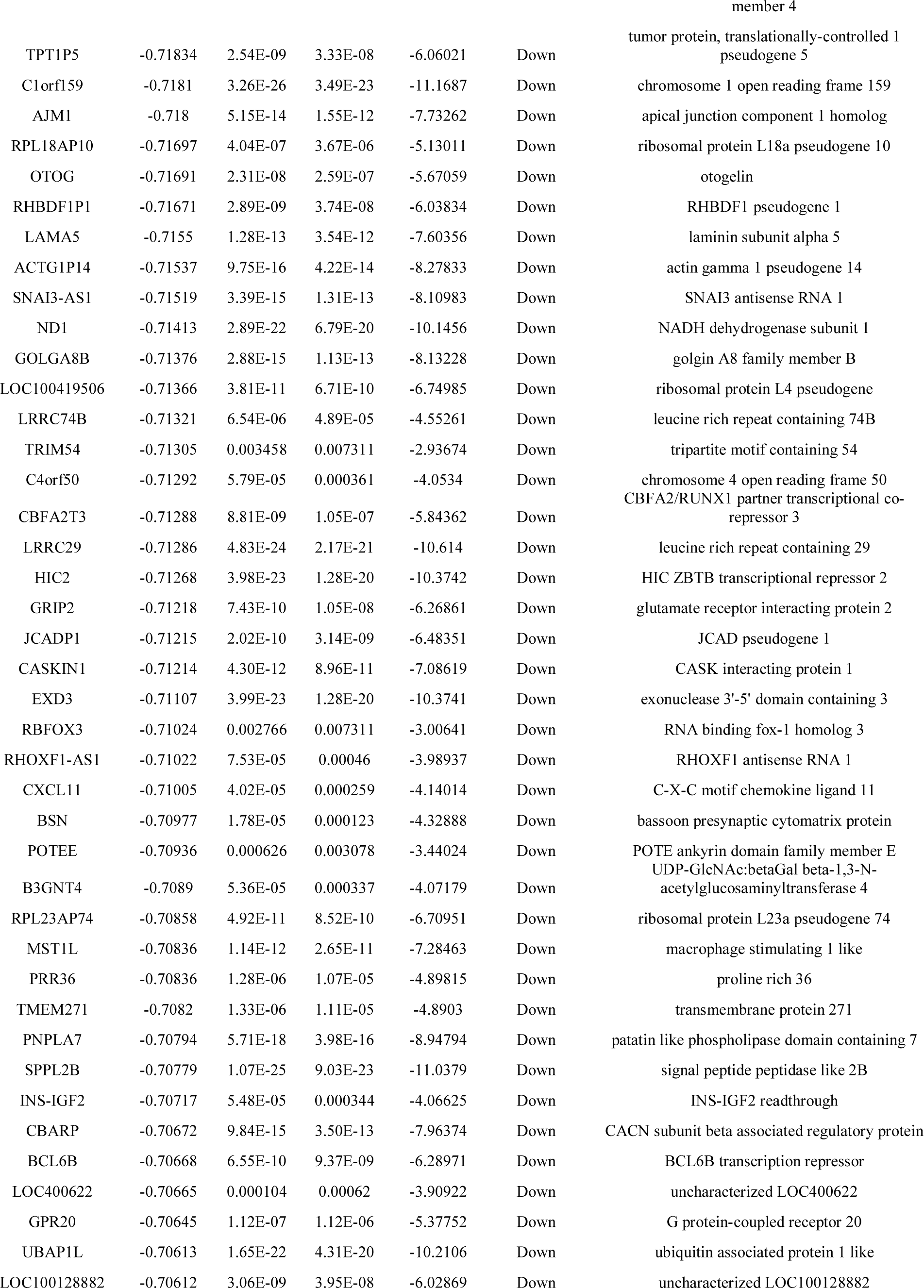
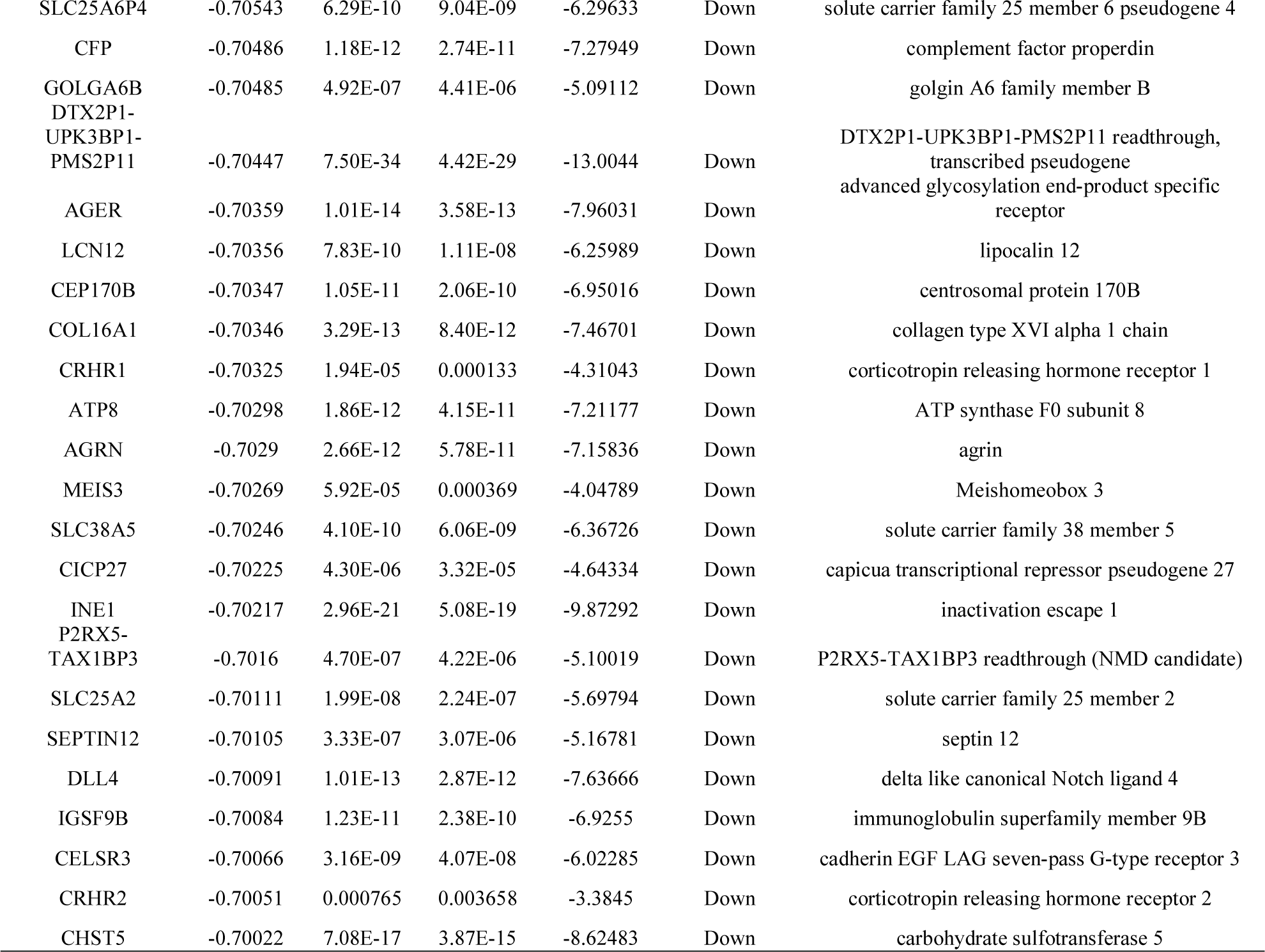
The statistical metrics for key differentially expressed genes (DEGs)

### GO and REACTOME pathway enrichment analysis

GO and REACTOME enrichment analysis for DEGs was performed using the ToppGene. We uploaded all DEGs to identify the GO categories and REACTOME pathways of the DEGs. The enriched GO terms were classified into biological process (BP), cellular component (CC) and molecular function (MF). GO analysis results showed that for BP, up regulated genes were significantly enriched in defense response and cell activation (Table 2). Down regulated genes were significantly enriched in BP, including ion transport and trans-synaptic signaling(Table 2). Up regulated genes were significantly enriched in CC, including whole membrane and intrinsic component of plasma membrane (Table 2). Down regulated genes were significantly enriched in CC, including integral component of plasma membrane and neuron projection (Table 2). GO MF showed that the up regulated genes were significantly enriched in molecular transducer activity and signaling receptor binding (Table 2). Down regulated genes were significantly enriched in MF, including transporter activity and transmembrane signaling receptor activity (Table 2). Moreover, REACTOME pathways were over represented in the up regulated genes, including neutrophil degranulation and innate immune system, while the down regulated genes were significantly enriched in REACTOME pathways, including neuronal system and NCAM signaling for neurite out-growth (Table 3).

**Table 2.**
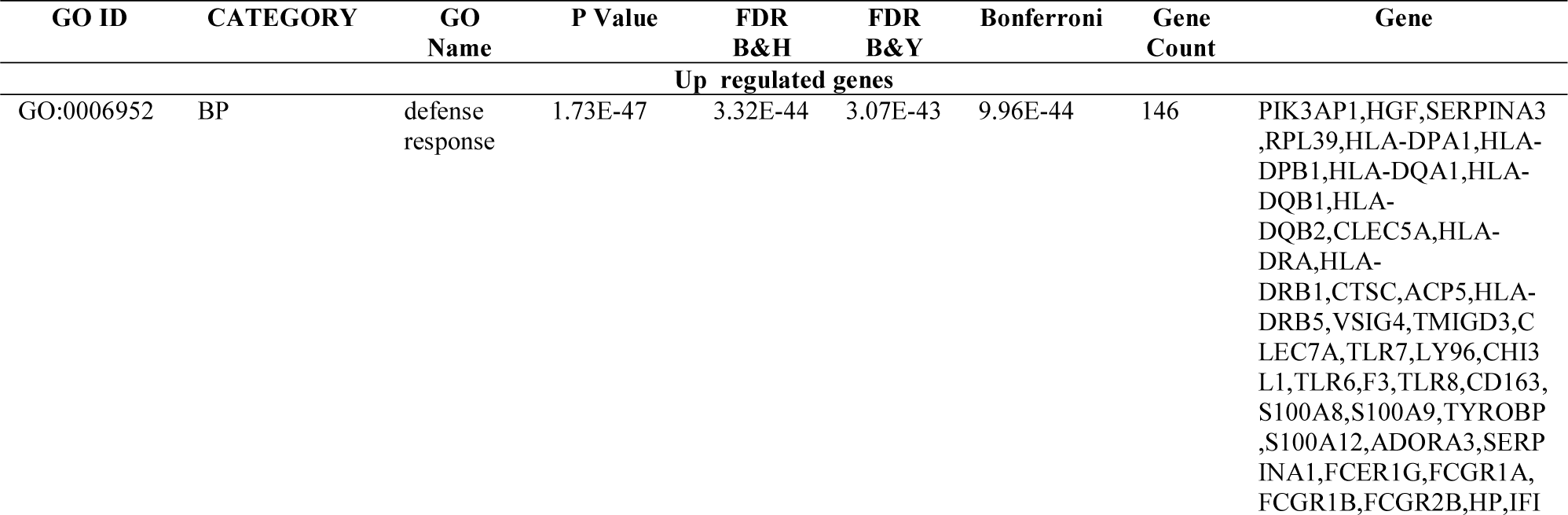
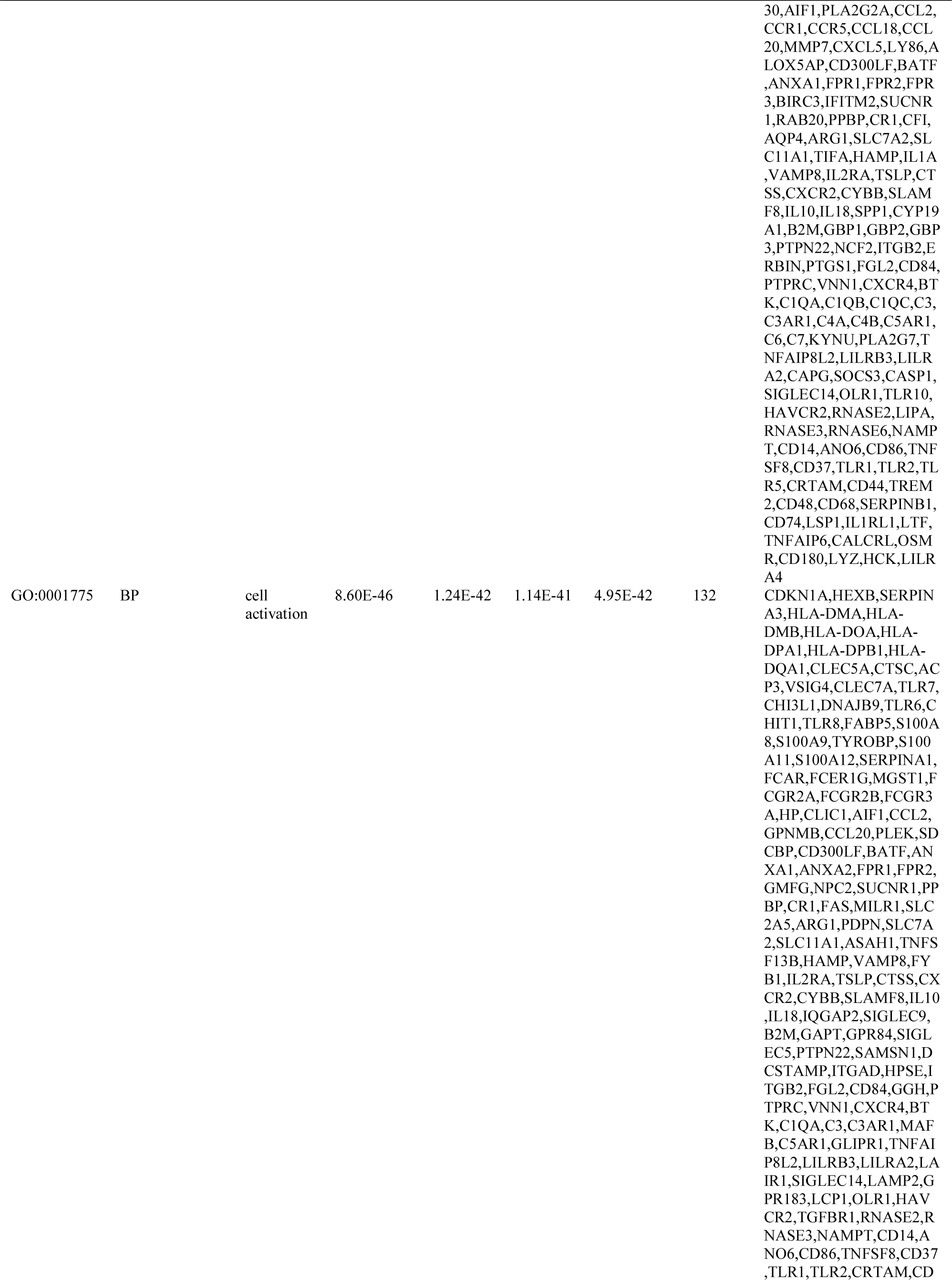
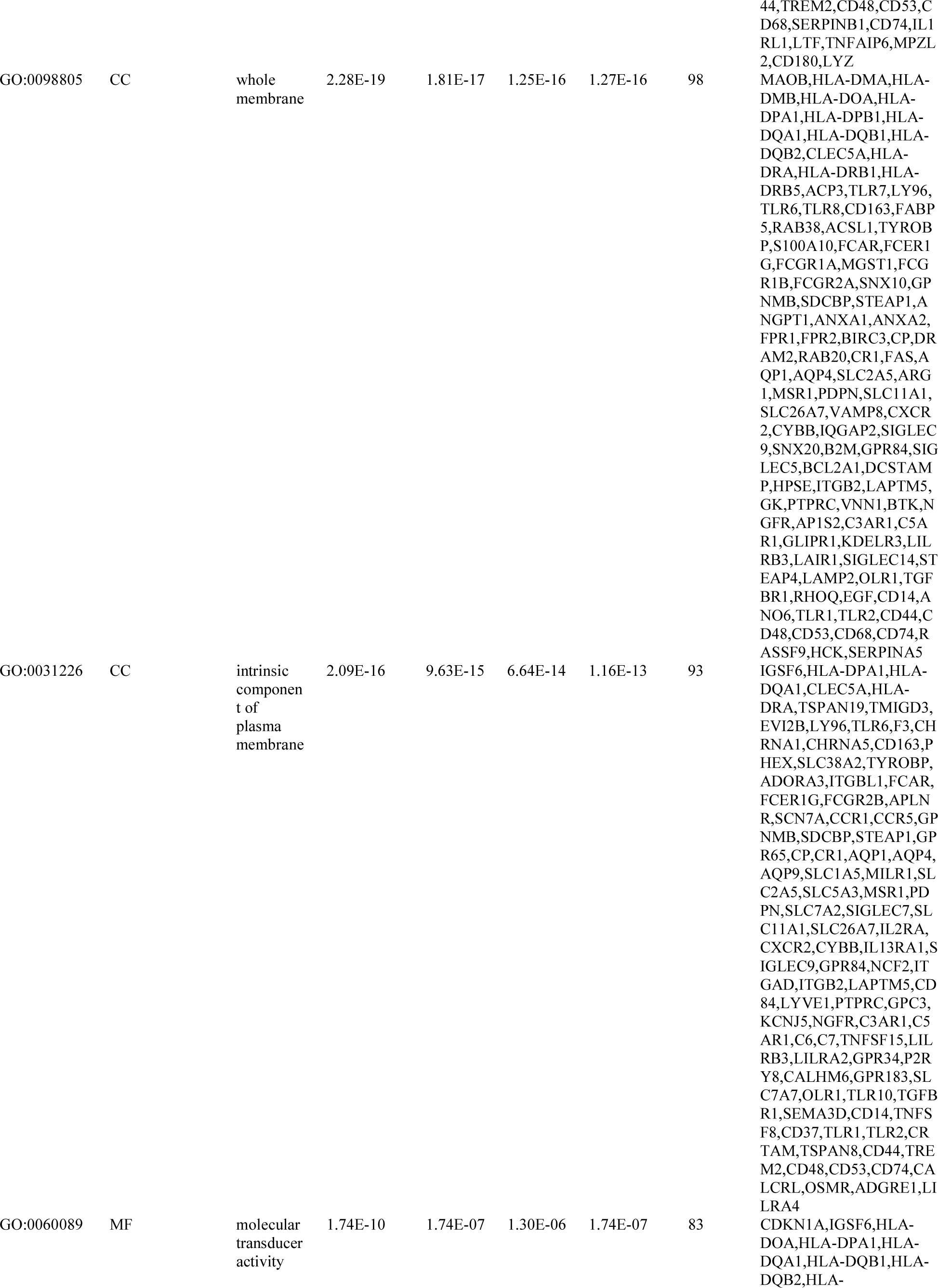
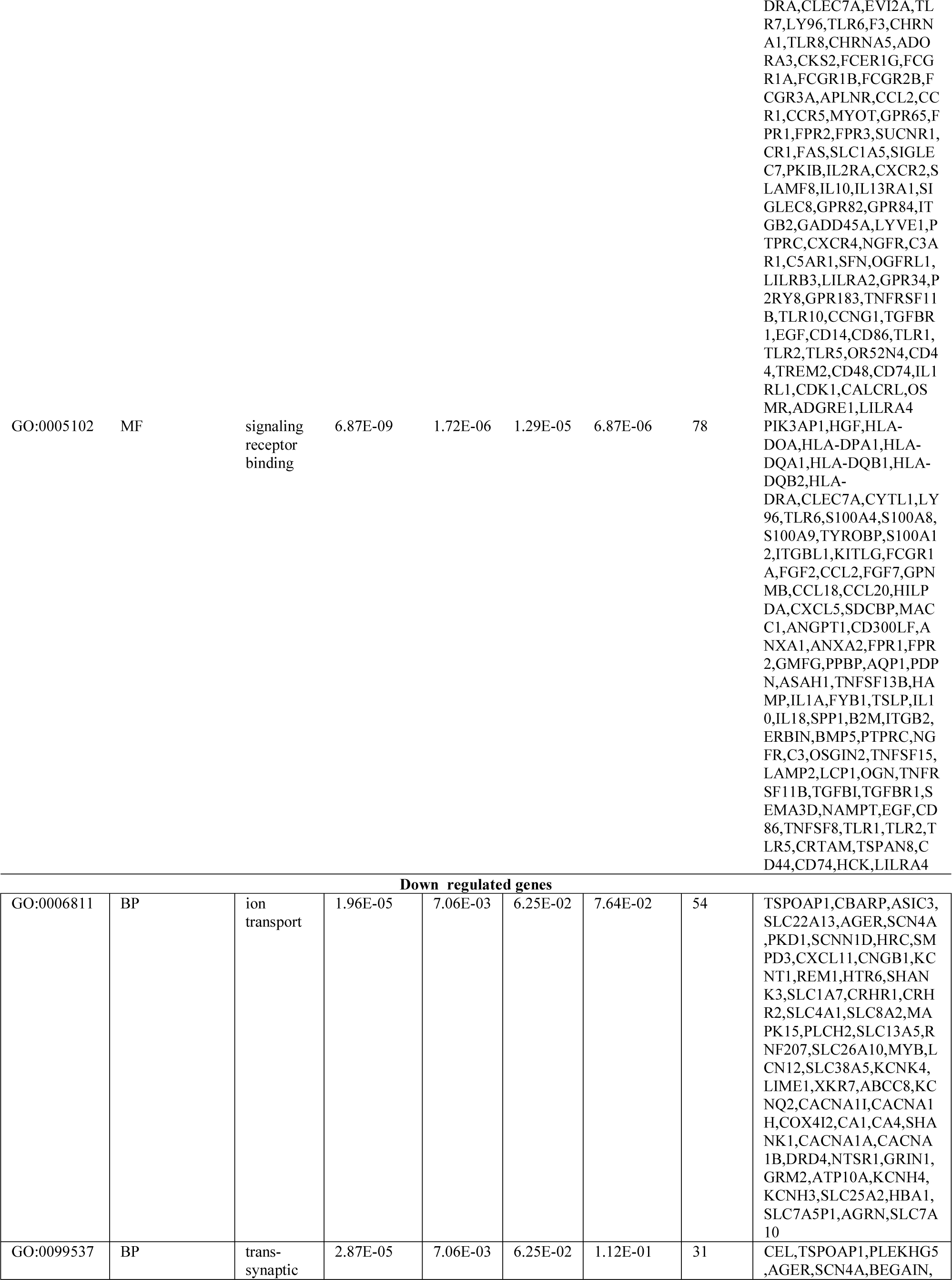
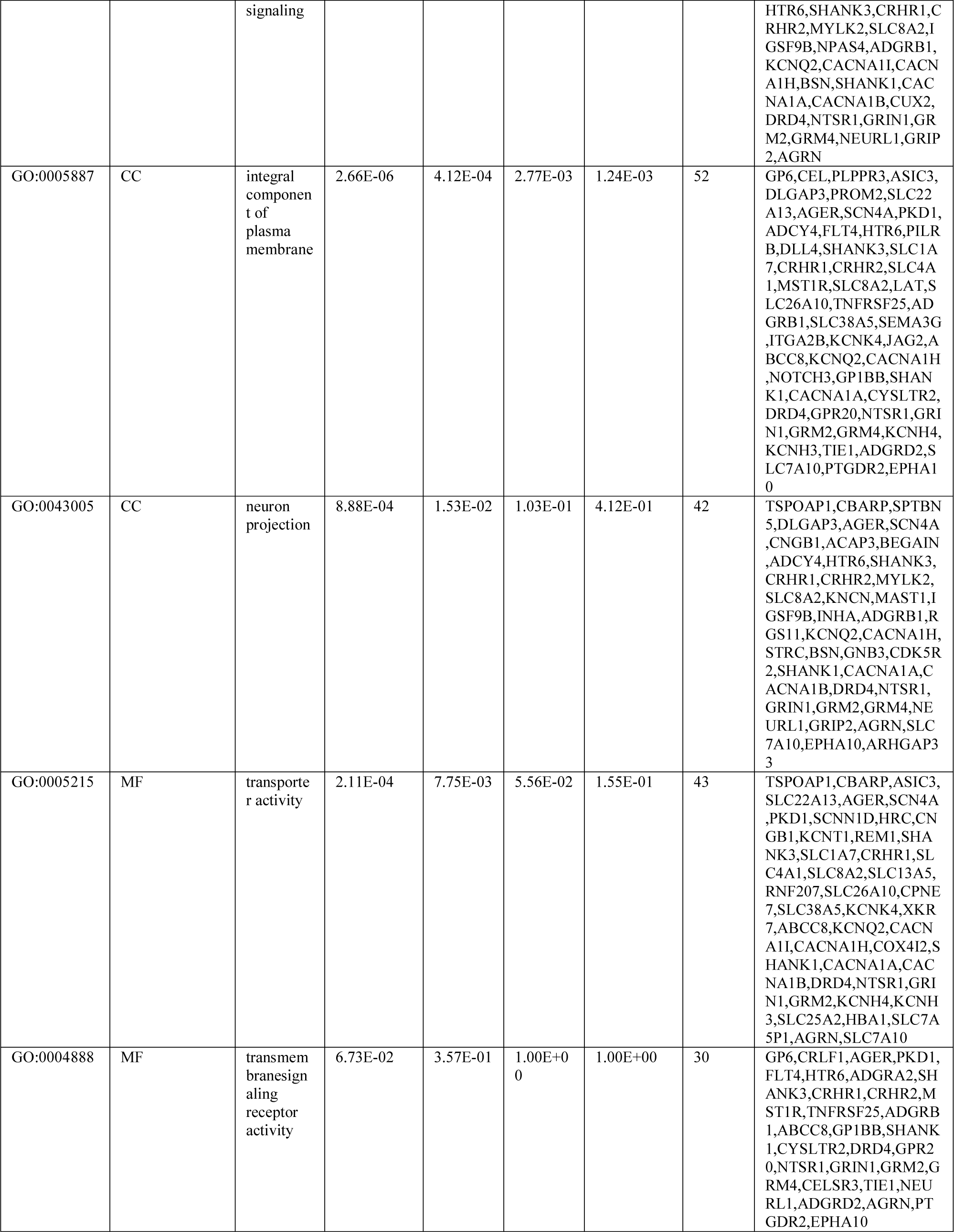
The enriched GO terms of the up and down regulated differentially expressed genes

**Table 3.** The enriched pathway terms of the up and down regulated differentially expressed genes

| Pathway ID | Pathway Name | P-value | FDR B&H | FDR B&Y | Bonferroni | Gene Count | Gene |
| --- | --- | --- | --- | --- | --- | --- | --- |
| 1457780 | Neutrophil degranulation | 8.98E-29 | 7.23E-26 | 5.26E-25 | 7.23E-26 | 67 | HEXB,SERPINA3,CLEC5A,CTSC,ACP3,CHI3L1,C<br>HIT1,FABP5,S100A8,S100A9,TYROBP,S100A11,S<br>100A12,SERPINA1,FCAR,FCER1G,MGST1,FCGR2<br>A,HP,SDCBP,ANXA2,FP<br>R1,FPR2,GMFG,NPC2,PP<br>BP,CR1,SLC2A5,ARG1,S<br>LC11A1,ASAH1,VAMP8,<br>CTSS,CXCR2,CYBB,IQG<br>AP2,SIGLEC9,B2M,GPR8<br>4,SIGLEC5,HPSE,ITGB2,<br>FGL2,GGH,PTPRC,VNN1<br>,C3,C3AR1,C5AR1,GLIP<br>R1,LILRB3,LAIR1,SIGLE<br>C14,LAMP2,OLR1,RNAS<br>E2,RNASE3,CD14,ANO6,<br>TLR2,CD44,CD53,CD68,S<br>ERPINB1,LTF,TNFAIP6,<br>LYZ |
| 1269203 | Innate Immune System | 4.32E-26 | 1.74E-23 | 1.27E-22 | 3.48E-23 | 106 | CDKN1A,HEXB,PIK3AP<br>1,HGF,SERPINA3,CLEC5<br>A,CTSC,ACP3,CLEC7A,T<br>LR7,LY96,CHI3L1,TLR6,<br>CHIT1,TLR8,FABP5,S100<br>A8,S100A9,TYROBP,S10<br>0A11,S100A12,SERPINA<br>1,FCAR,KITLG,FCER1G,<br>FCGR1A,MGST1,FCGR2<br>A,FCGR3A,HP,FGF2,PLA<br>2G2A,FGF7,LY86,SDCBP<br>,ANGPT1,ANXA2,FPR1,F<br>PR2,GMFG,BIRC3,NPC2,<br>PPBP,CR1,CFI,SLC2A5,A<br>RG1,SLC11A1,ASAH1,V<br>AMP8,IL2RA,CTSS,PRO<br>S1,CXCR2,CYBB,IQGAP<br>2,SIGLEC9,B2M,GPR84,S<br>IGLEC5,NCF2,HPSE,ITG<br>B2,FGL2,GGH,PTPRC,V<br>NN1,BTK,C1QA,C1QB,C<br>1QC,C3,C3AR1,C4A,C4B,<br>C5AR1,C6,C7,GLIPR1,LI<br>LRB3,LAIR1,CASP1,SIG<br>LEC14,LAMP2,OLR1,TL<br>R10,RNASE2,RNASE3,R<br>NASE6,EGF,CD14,ANO6,<br>CD86,TLR1,TLR2,TLR5,<br>CD44,TREM2,CD53,CD6<br>8,SERPINB1,LTF,TNFAI<br>P6,CD180,LYZ,HCK |
| 1269201 | Immunoregulatory interactions between a Lymphoid and a non-Lymphoid cell | 4.99E-09 | 3.66E-07 | 2.66E-06 | 4.02E-06 | 19 | TYROBP,FCGR1A,FCGR<br>2B,FCGR3A,CD300LF,LI<br>LRA6,SIGLEC7,SIGLEC9<br>,SIGLEC8,B2M,SIGLEC5,<br>ITGB2,C3,LILRB3,LILRA<br>2,LAIR1,CRTAM,TREM2<br>,LILRA4 |
| 1269576 | G alpha (i) signalling events | 3.24E-06 | 1.25E-04 | 9.05E-04 | 2.61E-03 | 21 | ADORA3,APLNR,CCR1,<br>CCR5,CCL20,CXCL5,AN<br>XA1,FPR1,FPR2,FPR3,SU<br>CNR1,PPBP,CXCR2,GNG<br>12,CXCR4,C3,C3AR1,C5<br>AR1,RGS1,RGS10,RGS18 |
| 1269318 | Signaling by Interleukins | 1.35E-04 | 4.18E-03 | 3.04E-02 | 1.09E-01 | 30 | CDKN1A,HGF,F13A1,KI<br>TLG,H3C11,H3C8,FGF2,<br>CCL2,FGF7,CCR1,CCR5,<br>CCL20,ANGPT1,BATF,A<br>NXA1,FPR1,IL1A,IL2RA,<br>IL10,IL13RA1,IL18,ITGB<br>2,SOCS3,CASP1,HAVCR<br>2,EGF,CD86,IL1RL1,OS<br>MR,HCK |
| 1269340 | Hemostasis | 2.78E-03 | 5.10E-02 | 3.71E-01 | 1.00E+00 | 30 | HGF,SERPINA3,F3,F5,F1<br>3A1,S100A10,SERPINA1,<br>FCER1G,H3C11,H3C8,PL<br>EK,ANGPT1,ANXA2,PPB<br>P,PDPN,SRGN,IL2RA,PR<br>OS1,ITGB2,CD84,GNG12<br>,LAMP2,SLC7A7,OLR1,K<br>IF20A,EGF,CD44,CD48,C<br>D74,SERPINA5 |
| 1269544 | GPCR ligand binding | 6.98E-03 | 1.00E-01 | 7.29E-01 | 1.00E+00 | 22 | ADORA3,APLNR,CCR1,<br>CCR5,CCL20,CXCL5,GP<br>R65,ANXA1,FPR1,FPR2,<br>FPR3,SUCNR1,PPBP,CX<br>CR2,GNG12,CXCR4,C3,C<br>3AR1,C5AR1,GPR183,CA<br>LCRL,ADGRE1 |
| **Down regulated genes** |  |  |  |  |  |  |  |
| 1268763 | Neuronal System | 1.07E-05 | 2.80E-03 | 1.91E-02 | 5.59E-03 | 17 | TSPOAP1,DLGAP3,BEG<br>AIN,ADCY4,SHANK3,SL<br>C1A7,KCNK4,ABCC8,KC<br>NQ2,GNB3,SHANK1,CA<br>CNA1A,CACNA1B,GRIN<br>1,KCNH4,KCNH3,GRIP2 |
| 1270312 | NCAM signaling for neurite out-growth | 6.61E-03 | 1.51E-01 | 1.00E+00 | 1.00E+00 | 10 | CNKSRI,SPTBN5,COL9<br>A1,LAT,ITGA2B,FGF17,<br>CACNA1I,CACNA1H,GR<br>IN1,AGRN |
| 1270303 | Axon guidance | 1.27E-02 | 1.81E-01 | 1.00E+00 | 1.00E+00 | 15 | CNKSRI,SPTBN5,SCN4A<br>,AGAP2,COL9A1,SHAN<br>K3,LAT,ITGA2B,FGF17,<br>KCNQ2,CACNA1I,CACN<br>A1H,GRIN1,AGRN,EPHA<br>10 |
| 1269544 | GPCR ligand binding | 1.32E-02 | 1.82E-01 | 1.00E+00 | 1.00E+00 | 13 | PLPPR3,CXCL11,HTR6,<br>WNT7B,CRHR1,CRHR2,<br>GNB3,CYSLTR2,DRD4,N<br>TSR1,GRM2,GRM4,PTG<br>DR2 |
| 1269903 | Transmembrane transport of small molecules | 1.05E-01 | 4.96E-01 | 1.00E+00 | 1.00E+00 | 14 | ASIC3,SCNN1D,ADCY4,<br>SLC1A7,SLC4A1,SLC8A<br>2,SLC13A5,LCN12,SLC3<br>8A5,LCN15,ABCC8,GNB<br>3,ATP10A,SLC7A10 |
| 1269340 | Hemostasis | 1.25E-01 | 4.96E-01 | 1.00E+00 | 1.00E+00 | 13 | GP6,SLC8A2,LAT,MYB,<br>KIF26A,KIF12,ITGA2B,I<br>TIH4,CABLES1,GNB3,G<br>P1BB,KIFC2,SLC7A10 |

### PPI network construction and module analysis

By using the STRING database, the PPI network of DEGs was established and consisted of 3837 nodes and 7530 edges (Fig .3). The PPI network was observed by Cytoscape, and the cut-off criterion of hub gene selection with high node degree, betweenness centrality, stress centrality and closeness centrality. A top 10 genes were selected for key biomarker identification (Table 4). They consisted of 5 up regulated genes (CDK1, TOP2A, MAD2L1, RSL24D1 and CDKN1A) and 5 down regulated genes (NOTCH3, MYB, PWP2, WNT7B and HSPA12B). The 2 most significant modules were extracted from the PPI network by PEWCC1, which included a module 1 consisted of 9 nodes and 17 edges (Fig.4A), which are mainly associated with molecular transducer activity and innate immune system and module 2 consisted of 8 nodes and 15 edges (Fig.4B), which are mainly associated with GPCR ligand binding, neuronal system, trans-synaptic signaling and integral component of plasma membrane.

**Fig. 3.**
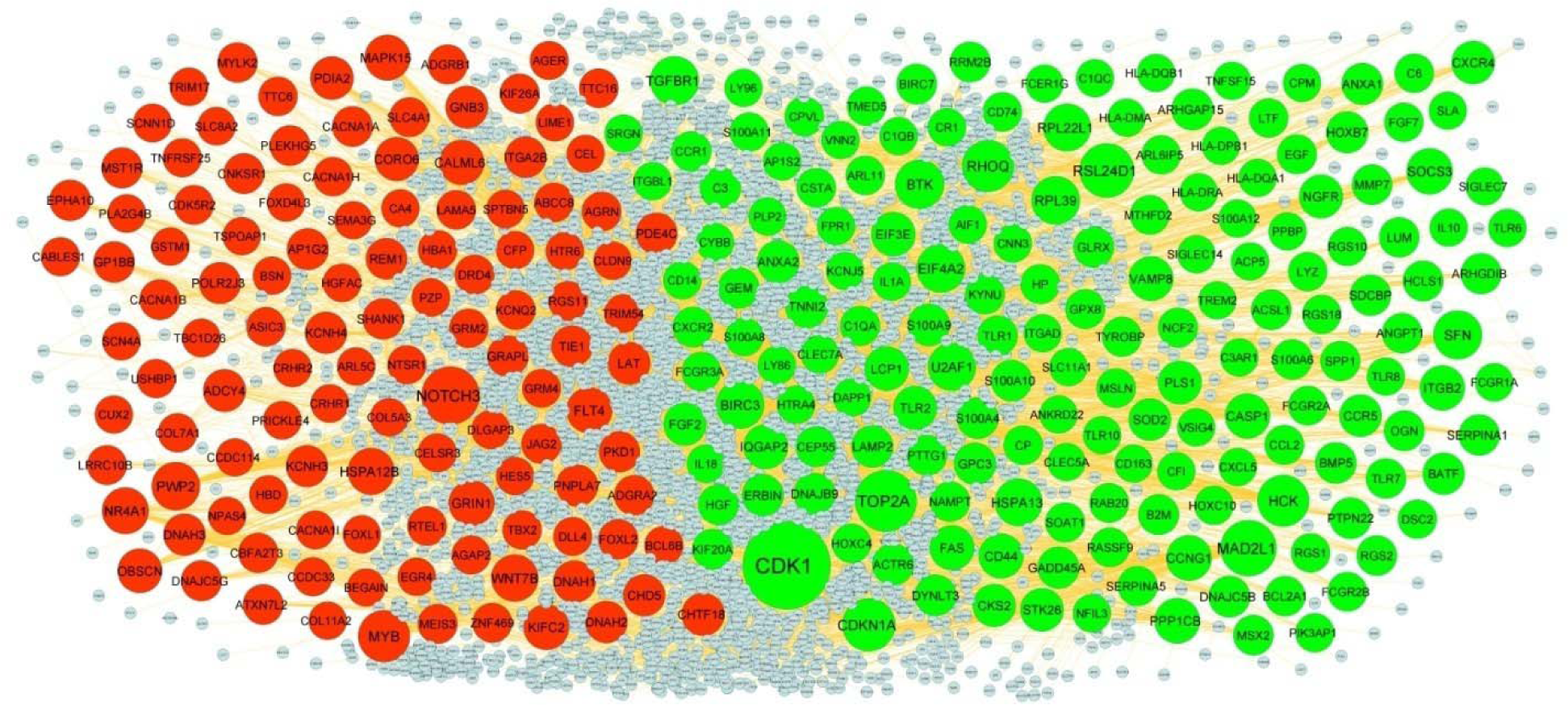
PPI network of DEGs. The PPI network of DEGs was constructed using Cytoscap. Up regulated genes are marked in green; down regulated genes are marked in red

**Fig. 4.**
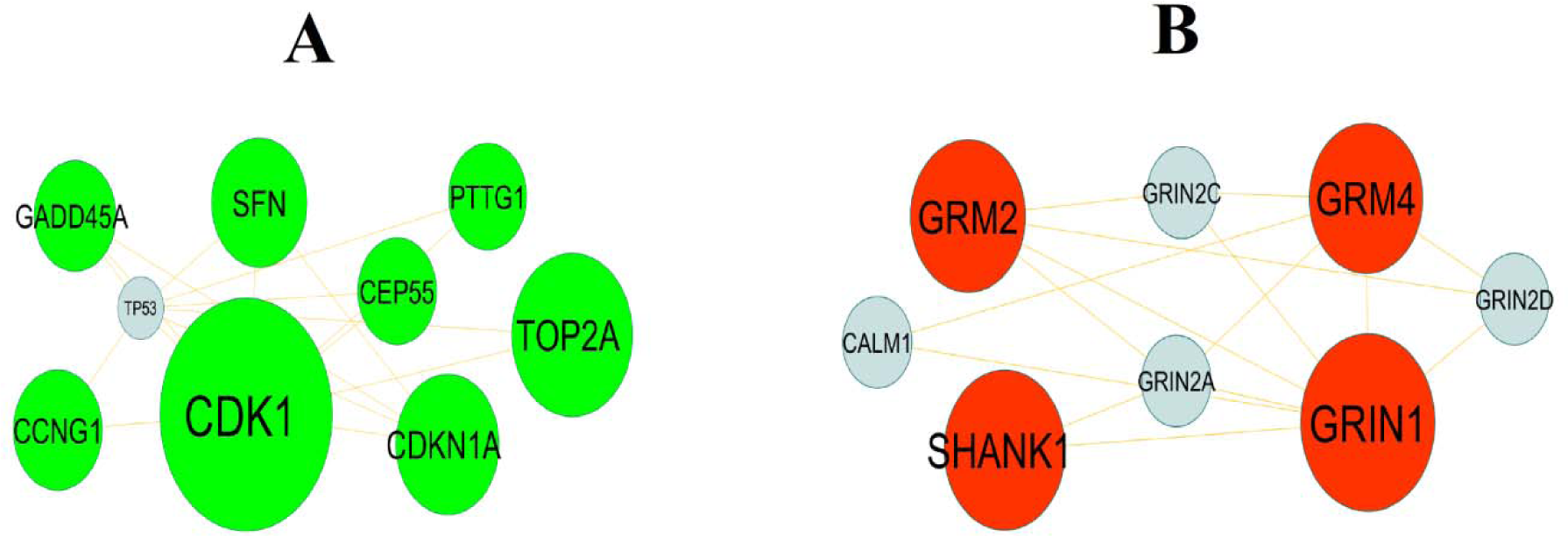
Modules of isolated form PPI of DEGs. (A) The most significant module was obtained from PPI network with 9 nodes and 17 edges for up regulated genes (B) The most significant module was obtained from PPI network with 8 nodes and 15 edges for down regulated genes. Up regulated genes are marked in green; down regulated genes are marked in red.

**Table 4.** Topology table for up and down regulated genes

| Category | Node | Degree | Betweenness | Stress | Closeness |
| --- | --- | --- | --- | --- | --- |
| Up | CDK1 | 402 | 0.197084 | 47028580 | 0.367116 |
| Up | TOP2A | 191 | 0.081627 | 13073002 | 0.341251 |
| Up | MAD2L1 | 146 | 0.040284 | 8144352 | 0.324233 |
| Up | RSL24D1 | 133 | 0.034596 | 8777264 | 0.30143 |
| Up | CDKN1A | 122 | 0.030348 | 6995294 | 0.331404 |
| Up | RHOQ | 111 | 0.052089 | 7439940 | 0.320307 |
| Up | HCK | 111 | 0.035255 | 11066684 | 0.322733 |
| Up | BTK | 90 | 0.02356 | 5280320 | 0.301359 |
| Up | TGFBR1 | 89 | 0.036693 | 10570252 | 0.318499 |
| Up | SFN | 88 | 0.030706 | 5847364 | 0.333362 |
| Up | EIF4A2 | 86 | 0.02546 | 4672448 | 0.306096 |
| Up | RPL39 | 85 | 0.014424 | 6331184 | 0.300297 |
| Up | BIRC3 | 82 | 0.02467 | 6593536 | 0.30953 |
| Up | U2AF1 | 80 | 0.03024 | 6114556 | 0.299524 |
| Up | HSPA13 | 79 | 0.023849 | 4111210 | 0.312225 |
| Up | RPL22L1 | 78 | 0.011772 | 4945670 | 0.300133 |
| Up | SOCS3 | 67 | 0.018611 | 4368764 | 0.312709 |
| Up | PPP1CB | 61 | 0.024464 | 3908348 | 0.30841 |
| Up | CCNG1 | 61 | 0.00591 | 1238240 | 0.315876 |
| Up | ITGB2 | 58 | 0.028427 | 5334096 | 0.316946 |
| Up | CASP1 | 58 | 0.018625 | 2471864 | 0.308981 |
| Up | CD44 | 56 | 0.025232 | 4779164 | 0.315098 |
| Up | VAMP8 | 54 | 0.02368 | 2898878 | 0.297872 |
| Up | IQGAP2 | 54 | 0.007736 | 2790576 | 0.313552 |
| Up | FAS | 54 | 0.014105 | 5340678 | 0.283707 |
| Up | EIF3E | 51 | 0.015213 | 3479548 | 0.297457 |
| Up | LCP1 | 51 | 0.009397 | 1985584 | 0.313347 |
| Up | NGFR | 47 | 0.016065 | 3746024 | 0.308459 |
| Up | TLR2 | 46 | 0.012377 | 2255708 | 0.298824 |
| Up | PLS1 | 46 | 0.007749 | 1638298 | 0.310834 |
| Up | STK26 | 44 | 0.017254 | 2794926 | 0.299664 |
| Up | CXCR4 | 44 | 0.017176 | 4261588 | 0.314607 |
| Up | TNNI2 | 42 | 0.00668 | 8295678 | 0.245961 |
| Up | CKS2 | 40 | 0.004636 | 968814 | 0.317892 |
| Up | FGF2 | 39 | 0.01534 | 1981338 | 0.289378 |
| Up | CCR5 | 37 | 0.008114 | 1662686 | 0.277388 |
| Up | ANXA2 | 37 | 0.014813 | 3649912 | 0.313963 |
| Up | SOD2 | 36 | 0.01232 | 2200890 | 0.297526 |
| Up | S100A9 | 36 | 0.008539 | 1250402 | 0.271902 |
| Up | GPC3 | 34 | 0.004643 | 1075058 | 0.239167 |
| Up | BATF | 33 | 0.00659 | 1842444 | 0.271191 |
| Up | HOXB7 | 32 | 0.005224 | 2604922 | 0.247644 |
| Up | ARHGAP15 | 32 | 0.006037 | 1575176 | 0.265688 |
| Up | GADD45A | 32 | 0.007594 | 1064172 | 0.31996 |
| Up | LAMP2 | 32 | 0.007823 | 3053920 | 0.254799 |
| Up | ACTR6 | 31 | 0.010083 | 2336460 | 0.236878 |
| Up | DNAJB9 | 31 | 0.002157 | 834952 | 0.264315 |
| Up | ANXA1 | 30 | 0.00939 | 2103520 | 0.301549 |
| Up | DNAJC5B | 30 | 0.001424 | 1100202 | 0.266278 |
| Up | PTPN22 | 30 | 0.003095 | 1739114 | 0.273025 |
| Up | ERBIN | 30 | 0.012972 | 2825982 | 0.312887 |
| Up | C3 | 29 | 0.011276 | 3367732 | 0.238335 |
| Up | S100A4 | 29 | 0.006124 | 1593532 | 0.303601 |
| Up | DYNLT3 | 29 | 0.004175 | 1869030 | 0.22383 |
| Up | BMP5 | 28 | 0.005191 | 833118 | 0.253134 |
| Up | GEM | 27 | 0.003998 | 3127856 | 0.244674 |
| Up | S100A10 | 27 | 0.010457 | 853366 | 0.299758 |
| Up | SDCBP | 26 | 0.007881 | 1442622 | 0.298081 |
| Up | SERPINA1 | 26 | 0.011803 | 1824802 | 0.301004 |
| Up | ARHGDIB | 25 | 0.00217 | 1032648 | 0.304275 |
| Up | GLRX | 23 | 0.006494 | 1256738 | 0.295623 |
| Up | B2M | 23 | 0.009862 | 1122844 | 0.292869 |
| Up | OGN | 22 | 0.003355 | 1546298 | 0.256554 |
| Up | HCLS1 | 22 | 0.004584 | 751498 | 0.2831 |
| Up | KIF20A | 22 | 0.002995 | 1053136 | 0.316946 |
| Up | BIRC7 | 21 | 0.002067 | 803284 | 0.295213 |
| Up | RRM2B | 21 | 0.004262 | 946508 | 0.293901 |
| Up | HGF | 20 | 0.006487 | 729002 | 0.264479 |
| Up | RGS2 | 20 | 0.003947 | 2415788 | 0.251971 |
| Up | MMP7 | 20 | 0.00531 | 450650 | 0.261611 |
| Up | BCL2A1 | 20 | 0.004104 | 382146 | 0.268816 |
| Up | MSX2 | 20 | 0.003782 | 1705198 | 0.231098 |
| Up | CXCR2 | 20 | 0.005673 | 1759130 | 0.230723 |
| Up | PTTG1 | 20 | 0.003325 | 436594 | 0.313552 |
| Up | CEP55 | 20 | 0.005756 | 585852 | 0.31559 |
| Up | ACSL1 | 19 | 0.008972 | 975302 | 0.291512 |
| Up | LUM | 19 | 0.001751 | 1357886 | 0.250245 |
| Up | TLR7 | 19 | 0.002068 | 466520 | 0.26195 |
| Up | CCR1 | 18 | 0.002536 | 1830672 | 0.225753 |
| Up | TLR1 | 18 | 1.00E-03 | 270250 | 0.25409 |
| Up | S100A6 | 18 | 0.005993 | 566808 | 0.300274 |
| Up | ANGPT1 | 18 | 0.001995 | 247354 | 0.260297 |
| Up | NAMPT | 17 | 0.006404 | 866924 | 0.292757 |
| Up | TYROBP | 16 | 0.006739 | 3350394 | 0.21767 |
| Up | SERPINA5 | 16 | 0.005047 | 1140526 | 0.204445 |
| Up | SOAT1 | 16 | 0.007372 | 834918 | 0.293183 |
| Up | C1QA | 16 | 0.003569 | 907138 | 0.244051 |
| Up | SPP1 | 16 | 0.003897 | 481758 | 0.273668 |
| Up | EGF | 15 | 0.003463 | 1877948 | 0.246736 |
| Up | NCF2 | 15 | 0.003176 | 846020 | 0.252984 |
| Up | GPX8 | 15 | 0.003073 | 745356 | 0.29249 |
| Up | CCL2 | 14 | 0.002658 | 610374 | 0.238379 |
| Up | CPVL | 14 | 0.004372 | 636758 | 0.294624 |
| Up | RAB20 | 14 | 0.006334 | 466976 | 0.201863 |
| Up | KYNU | 14 | 0.004584 | 733080 | 0.292534 |
| Up | HP | 14 | 0.005264 | 634858 | 0.244051 |
| Up | MTHFD2 | 14 | 0.006469 | 723768 | 0.291556 |
| Up | FCGR2A | 13 | 0.002775 | 672196 | 0.305195 |
| Up | FCGR2B | 13 | 0.002454 | 818192 | 0.253118 |
| Up | RASSF9 | 13 | 0.003768 | 327690 | 0.246181 |
| Up | CD74 | 12 | 0.00413 | 277206 | 0.30281 |
| Up | CSTA | 12 | 0.00461 | 539282 | 0.291489 |
| Up | CP | 12 | 0.0037 | 233612 | 0.236849 |
| Up | MSLN | 12 | 0.004721 | 1874812 | 0.200764 |
| Up | LTF | 11 | 0.006533 | 917294 | 0.301099 |
| Up | C6 | 11 | 0.002184 | 776286 | 0.231461 |
| Up | ARL11 | 11 | 0.003483 | 486670 | 0.292869 |
| Up | TMED5 | 11 | 0.003557 | 612564 | 0.29672 |
| Up | LYZ | 10 | 0.002369 | 389198 | 0.296079 |
| Up | ARL6IP5 | 10 | 0.004541 | 404738 | 0.297249 |
| Up | FGF7 | 10 | 0.001158 | 492864 | 0.211408 |
| Up | HOXC4 | 10 | 0.001703 | 539558 | 0.220917 |
| Up | HOXC10 | 10 | 0.001805 | 548562 | 0.216662 |
| Up | FCGR3A | 10 | 0.001167 | 993240 | 0.224814 |
| Up | RGS1 | 10 | 1.49E-04 | 62882 | 0.218265 |
| Up | S100A11 | 9 | 0.001746 | 228728 | 0.296354 |
| Up | DSC2 | 9 | 0.002704 | 467128 | 0.222106 |
| Up | PIK3AP1 | 9 | 0.001407 | 319970 | 0.250147 |
| Up | RGS18 | 9 | 1.22E-04 | 135174 | 0.215457 |
| Up | RGS10 | 9 | 0.002616 | 1549108 | 0.303649 |
| Up | S100A8 | 3 | 6.23E-05 | 17078 | 0.23881 |
| Up | FCGR1A | 3 | 8.90E-05 | 110248 | 0.245034 |
| Up | CD14 | 3 | 1.41E-04 | 20250 | 0.249886 |
| Up | TLR10 | 2 | 0 | 0 | 0.230418 |
| Up | CR1 | 2 | 7.30E-05 | 24248 | 0.202417 |
| Up | CNN3 | 2 | 1.30E-07 | 104 | 0.238721 |
| Up | TLR6 | 2 | 0 | 0 | 0.238929 |
| Up | SLC11A1 | 1 | 0 | 0 | 0.191503 |
| Up | TLR8 | 1 | 0 | 0 | 0.231587 |
| Up | CXCL5 | 1 | 0 | 0 | 0.187479 |
| Up | PPBP | 1 | 0 | 0 | 0.187479 |
| Up | SIGLEC7 | 1 | 0 | 0 | 0.238231 |
| Up | FCER1G | 1 | 0 | 0 | 0.183558 |
| Up | C1QB | 1 | 0 | 0 | 0.196185 |
| Up | C1QC | 1 | 0 | 0 | 0.196185 |
| Up | S100A12 | 1 | 0 | 0 | 0.197508 |
| Up | TNFSF15 | 1 | 0 | 0 | 0.191647 |
| Up | LY86 | 1 | 0 | 0 | 0.207587 |
| Up | ACP5 | 1 | 0 | 0 | 0.214878 |
| Up | SRGN | 1 | 0 | 0 | 0.239615 |
| Up | IL1A | 1 | 0 | 0 | 0.236062 |
| Up | IL18 | 1 | 0 | 0 | 0.236062 |
| Up | LY96 | 1 | 0 | 0 | 0.230086 |
| Up | NFIL3 | 1 | 0 | 0 | 0.213348 |
| Up | C3AR1 | 1 | 0 | 0 | 0.192474 |
| Up | CLEC7A | 1 | 0 | 0 | 0.230086 |
| Up | FPR1 | 1 | 0 | 0 | 0.231698 |
| Up | HTRA4 | 1 | 0 | 0 | 0.236382 |
| Up | VNN2 | 1 | 0 | 0 | 0.240683 |
| Up | AP1S2 | 1 | 0 | 0 | 0.228075 |
| Up | CPM | 1 | 0 | 0 | 0.197916 |
| Up | HLA-DQA1 | 1 | 0 | 0 | 0.232443 |
| Up | KCNJ5 | 1 | 0 | 0 | 0.187736 |
| Up | CD163 | 1 | 0 | 0 | 0.196185 |
| Up | SIGLEC14 | 1 | 0 | 0 | 0.178768 |
| Up | ANKRD22 | 1 | 0 | 0 | 0.218029 |
| Up | TREM2 | 1 | 0 | 0 | 0.178768 |
| Up | VSIG4 | 1 | 0 | 0 | 0.192474 |
| Up | HLA-DMA | 1 | 0 | 0 | 0.232443 |
| Up | AIF1 | 1 | 0 | 0 | 0.238602 |
| Up | ITGBL1 | 1 | 0 | 0 | 0.212615 |
| Up | PLP2 | 1 | 0 | 0 | 0.184184 |
| Up | CYBB | 1 | 0 | 0 | 0.201916 |
| Up | ITGAD | 1 | 0 | 0 | 0.240683 |
| Up | CFI | 1 | 0 | 0 | 0.192474 |
| Up | HLA-DRA | 1 | 0 | 0 | 0.232443 |
| Up | HLA-DQB1 | 1 | 0 | 0 | 0.232443 |
| Up | SLA | 1 | 0 | 0 | 0.23226 |
| Up | HLA-DPB1 | 1 | 0 | 0 | 0.232443 |
| Up | IL10 | 1 | 0 | 0 | 0.230086 |
| Up | DAPP1 | 1 | 0 | 0 | 0.231587 |
| Up | CLEC5A | 1 | 0 | 0 | 0.178768 |
| Down | NOTCH3 | 164 | 0.091402 | 23838154 | 0.331777 |
| Down | MYB | 114 | 0.041442 | 7392588 | 0.330405 |
| Down | PWP2 | 85 | 0.032395 | 5057778 | 0.296812 |
| Down | WNT7B | 76 | 0.019782 | 2626426 | 0.277709 |
| Down | HSPA12B | 73 | 0.006667 | 2161072 | 0.27609 |
| Down | NR4A1 | 72 | 0.025505 | 4889526 | 0.31345 |
| Down | FLT4 | 66 | 0.020378 | 5735870 | 0.321085 |
| Down | MAPK15 | 64 | 0.021292 | 3346718 | 0.301643 |
| Down | GRIN1 | 64 | 0.023937 | 4117674 | 0.295965 |
| Down | KIFC2 | 63 | 0.016898 | 4559514 | 0.286568 |
| Down | CALML6 | 59 | 0.009409 | 2030892 | 0.270981 |
| Down | OBSCN | 54 | 0.008442 | 8694380 | 0.248188 |
| Down | CORO6 | 51 | 0.005529 | 1012934 | 0.270961 |
| Down | PDIA2 | 45 | 0.012294 | 5298738 | 0.267821 |
| Down | CHTF18 | 43 | 0.008614 | 1782478 | 0.30522 |
| Down | ADCY4 | 41 | 0.011029 | 6899344 | 0.256743 |
| Down | GNB3 | 40 | 0.010425 | 6470774 | 0.261736 |
| Down | LAT | 39 | 0.009974 | 2962082 | 0.3025 |
| Down | DNAH1 | 38 | 0.002688 | 5473740 | 0.231489 |
| Down | DNAH3 | 35 | 0.002569 | 5296504 | 0.231447 |
| Down | DNAH2 | 35 | 0.002776 | 5189040 | 0.232584 |
| Down | KCNH3 | 34 | 0.007586 | 1205362 | 0.212568 |
| Down | CHD5 | 34 | 0.009884 | 1545622 | 0.296537 |
| Down | KCNH4 | 34 | 0.007586 | 1205362 | 0.212568 |
| Down | PNPLA7 | 34 | 0.008145 | 3168532 | 0.244191 |
| Down | AP1G2 | 33 | 0.014151 | 1680394 | 0.295441 |
| Down | POLR2J3 | 33 | 0.015995 | 1968346 | 0.294127 |
| Down | PDE4C | 31 | 0.005427 | 2684966 | 0.25 |
| Down | TIE1 | 30 | 0.003319 | 1939470 | 0.246545 |
| Down | REM1 | 29 | 0.004314 | 1853664 | 0.270179 |
| Down | ITGA2B | 29 | 0.008146 | 945330 | 0.270008 |
| Down | EPHA10 | 29 | 0.00243 | 2289996 | 0.235742 |
| Down | GP1BB | 29 | 0.003547 | 2615204 | 0.260403 |
| Down | GRAPL | 29 | 0.005118 | 2545348 | 0.25409 |
| Down | SHANK1 | 28 | 0.009431 | 998326 | 0.259119 |
| Down | PKD1 | 26 | 0.00671 | 1877922 | 0.302071 |
| Down | MST1R | 26 | 0.005428 | 1662648 | 0.310658 |
| Down | DNAJC5G | 25 | 4.01E-04 | 267002 | 0.263425 |
| Down | CACNA1A | 23 | 0.008785 | 1982682 | 0.30143 |
| Down | CNKSR1 | 23 | 0.006291 | 3201818 | 0.241927 |
| Down | PLEKHG5 | 23 | 0.00133 | 464342 | 0.26406 |
| Down | CACNA1B | 23 | 0.010784 | 1739420 | 0.301786 |
| Down | ATXN7L2 | 22 | 0.009932 | 1793842 | 0.20617 |
| Down | MYLK2 | 20 | 0.00364 | 1889536 | 0.254174 |
| Down | ADGRA2 | 20 | 0.002251 | 1605348 | 0.251162 |
| Down | JAG2 | 19 | 0.003135 | 341490 | 0.270789 |
| Down | FOXL2 | 19 | 0.004964 | 2129576 | 0.251277 |
| Down | CBFA2T3 | 18 | 0.005802 | 2970336 | 0.235496 |
| Down | DRD4 | 17 | 0.005015 | 2287930 | 0.231112 |
| Down | ADGRB1 | 17 | 0.007025 | 1023980 | 0.296125 |
| Down | SLC4A1 | 16 | 0.005579 | 1584818 | 0.249528 |
| Down | CDK5R2 | 16 | 0.001136 | 151486 | 0.279531 |
| Down | RGS11 | 16 | 0.002225 | 1716534 | 0.214974 |
| Down | KIF26A | 16 | 0.002655 | 1291994 | 0.248462 |
| Down | LRRC10B | 16 | 5.10E-04 | 746332 | 0.248945 |
| Down | ZNF469 | 16 | 0.005113 | 850154 | 0.221172 |
| Down | MEIS3 | 16 | 0.003919 | 544578 | 0.252784 |
| Down | AGRN | 15 | 0.003672 | 694362 | 0.296262 |
| Down | LAMA5 | 14 | 0.004736 | 1018384 | 0.211641 |
| Down | RTEL1 | 14 | 0.005158 | 701906 | 0.292267 |
| Down | KCNQ2 | 14 | 0.012494 | 2580574 | 0.244565 |
| Down | ABCC8 | 14 | 0.003882 | 1444224 | 0.250653 |
| Down | TBX2 | 13 | 0.002103 | 795172 | 0.237582 |
| Down | HBA1 | 13 | 0.00606 | 870296 | 0.232217 |
| Down | TTC16 | 13 | 0.002938 | 2785128 | 0.239825 |
| Down | ASIC3 | 12 | 0.003761 | 656668 | 0.241805 |
| Down | CACNA1H | 12 | 0.002209 | 244700 | 0.238587 |
| Down | COL7A1 | 12 | 0.003033 | 1172566 | 0.20475 |
| Down | GRM2 | 12 | 0.002335 | 349216 | 0.236805 |
| Down | TTC6 | 12 | 0.002417 | 2559988 | 0.239795 |
| Down | USHBP1 | 10 | 0.002714 | 449390 | 0.221363 |
| Down | CABLES1 | 10 | 5.91E-04 | 469160 | 0.246039 |
| Down | BCL6B | 10 | 0.003553 | 579104 | 0.292892 |
| Down | NTSR1 | 10 | 0.002558 | 556770 | 0.216381 |
| Down | CCDC33 | 10 | 0.005748 | 1111714 | 0.203404 |
| Down | CELSR3 | 9 | 2.04E-04 | 146192 | 0.201672 |
| Down | DLL4 | 9 | 5.96E-04 | 69464 | 0.263281 |
| Down | DLGAP3 | 9 | 5.84E-04 | 140240 | 0.230987 |
| Down | TNFRSF25 | 9 | 0.001238 | 364564 | 0.237068 |
| Down | AGAP2 | 9 | 0.001386 | 378968 | 0.226273 |
| Down | AGER | 8 | 0.001241 | 255066 | 0.246103 |
| Down | GRM4 | 8 | 4.80E-04 | 146670 | 0.242233 |
| Down | SCNN1D | 8 | 0.001816 | 281882 | 0.291025 |
| Down | CACNA1I | 8 | 0.001429 | 289364 | 0.235742 |
| Down | PLA2G4B | 7 | 0.001088 | 429840 | 0.241958 |
| Down | TBC1D26 | 7 | 0.003666 | 372828 | 0.215494 |
| Down | TRIM17 | 6 | 8.96E-04 | 142824 | 0.292936 |
| Down | PZP | 6 | 7.80E-04 | 420298 | 0.199979 |
| Down | TRIM54 | 6 | 0.001716 | 358752 | 0.219979 |
| Down | PRICKLE4 | 6 | 4.54E-04 | 135962 | 0.225475 |
| Down | HBD | 6 | 0.0021 | 332484 | 0.188557 |
| Down | HGFAC | 6 | 0.001463 | 255144 | 0.219125 |
| Down | SLC8A2 | 5 | 4.48E-05 | 11786 | 0.231517 |
| Down | SPTBN5 | 5 | 5.71E-04 | 188288 | 0.207744 |
| Down | FOXD4L3 | 5 | 0.001565 | 1879774 | 0.173066 |
| Down | ARL5C | 5 | 4.48E-05 | 10724 | 0.201746 |
| Down | CRHR1 | 5 | 0.002615 | 1085972 | 0.202813 |
| Down | COL5A3 | 4 | 5.51E-04 | 233718 | 0.207295 |
| Down | LIME1 | 4 | 2.25E-05 | 12626 | 0.240215 |
| Down | GSTM1 | 4 | 0.001073 | 135528 | 0.291578 |
| Down | TSPOAP1 | 4 | 6.75E-04 | 133888 | 0.215264 |
| Down | HTR6 | 4 | 2.33E-04 | 111870 | 0.235511 |
| Down | CRHR2 | 4 | 5.22E-04 | 358806 | 0.144352 |
| Down | BSN | 4 | 0.001564 | 139830 | 0.188344 |
| Down | CLDN9 | 4 | 5.31E-04 | 151246 | 0.20146 |
| Down | SCN4A | 4 | 2.58E-04 | 89070 | 0.21238 |
| Down | EGR4 | 4 | 0.001564 | 182706 | 0.207363 |
| Down | SEMA3G | 3 | 6.53E-04 | 157670 | 0.203491 |
| Down | CUX2 | 3 | 1.12E-04 | 22356 | 0.23996 |
| Down | CCDC114 | 3 | 0.001043 | 2377910 | 0.15888 |
| Down | NPAS4 | 3 | 0.001043 | 201686 | 0.144651 |
| Down | CFP | 3 | 2.55E-04 | 58480 | 0.227656 |
| Down | FOXL1 | 3 | 6.86E-06 | 714 | 0.203642 |
| Down | BEGAIN | 3 | 0.001043 | 140566 | 0.217448 |
| Down | COL11A2 | 3 | 5.22E-04 | 200148 | 0.20335 |
| Down | HES5 | 3 | 2.07E-05 | 8016 | 0.224446 |
| Down | CA4 | 3 | 0.001043 | 360206 | 0.19975 |
| Down | CEL | 1 | 0 | 0 | 0.231433 |

### MiRNA-hub gene regulatory network construction

The miRNA-hub gene regulatory network was shown in Fig. 5, consisting of 2303 nodes and 9686 interactions. The nodes with high topological score can be regarded as key network nodes. The results, output by the powerful visualization system, indicated that 444 miRNAs (ex; hsa-mir-1248) might target CDKN1A, 127 miRNAs (ex; hsa-mir-6887-3p) might target TGFBR1, 113 miRNAs (ex; hsa-mir-101-5p) might target EIF4A2, 109 miRNAs (ex; hsa-mir-488-5p) might target CDK1, 107 miRNAs (ex; hsa-mir-548l) might target HSPA13, 88 miRNAs (ex; hsa-mir-6886-5p) might target WNT7B, 56 miRNAs (ex; hsa-mir-107) might target CHTF18, 47 miRNAs (ex; hsa-mir-150-3p) might target MYB, 43 miRNAs (ex; hsa-mir-195-5p) might target NR4A1 and 41 miRNAs (ex; hsa-mir-206) might target NOTCH3 and are listed in Table 5.

**Fig. 5.**
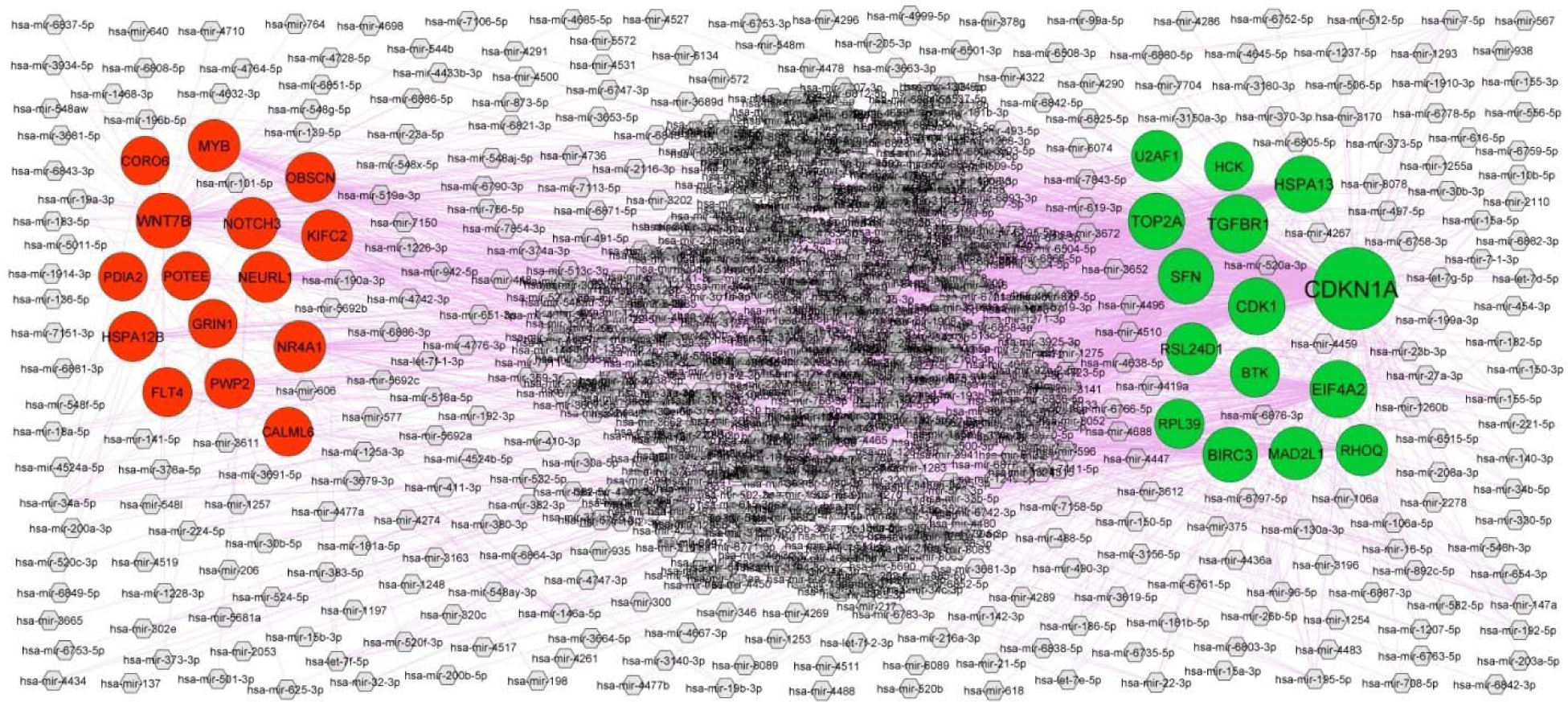
MiRNA - hub gene regulatory network. The purple color diamond nodes represent the key miRNAs; up regulated genes are marked in green; down regulated genes are marked in red.

**Table 5.**
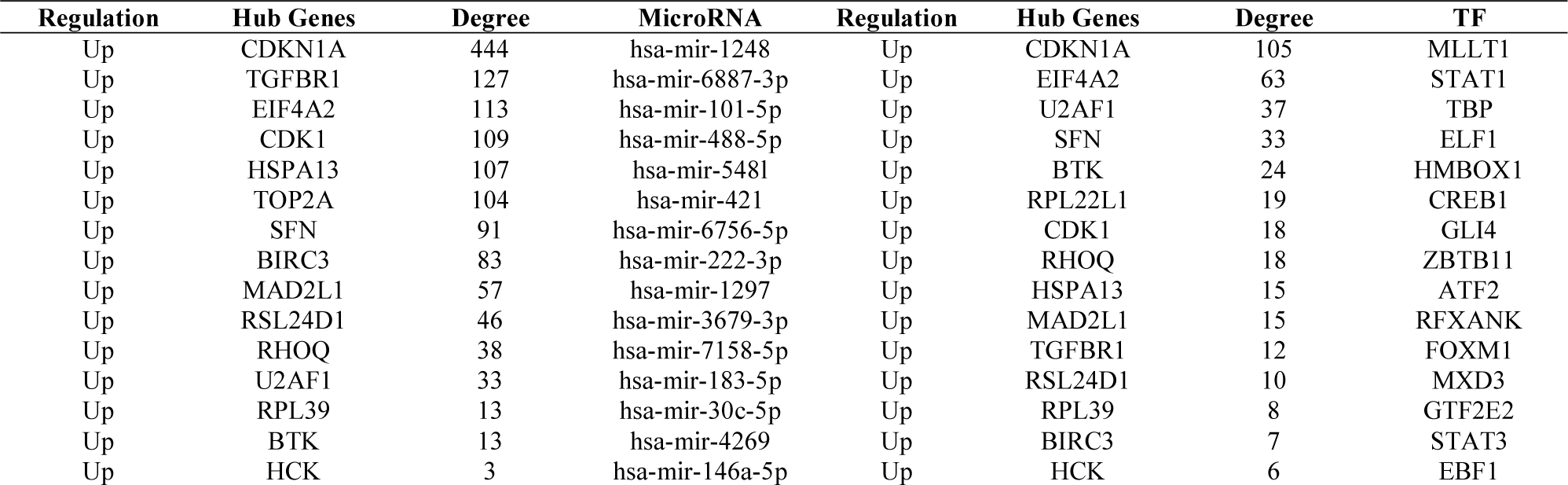
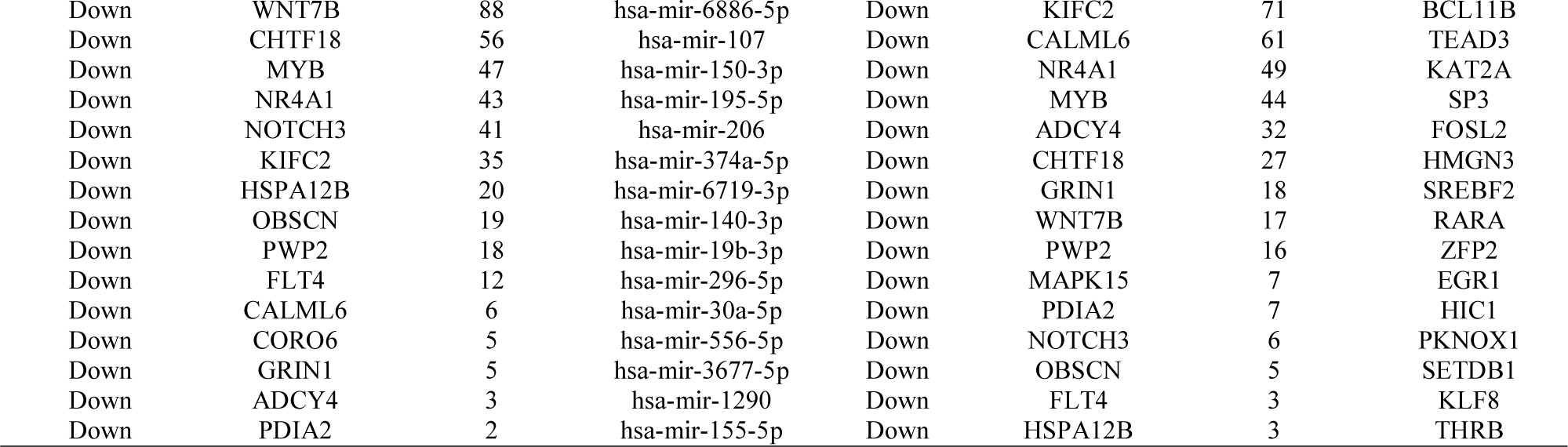
miRNA - hub gene and TF – hub gene interaction

### TF-hub gene regulatory network construction

The TF-hub gene regulatory network was shown in Fig. 6, consisting of 577 nodes and 5405 interactions. The nodes with high topological score can be regarded as key network nodes. The results, output by the powerful visualization system, indicated that 105 TFs (ex; MLLT1) might target CDKN1A, 63 TFs (ex; STAT1) might target EIF4A2, 37 TFs (ex; TBP) might target U2AF1, 33 TFs (ex; ELF1) might target SFN, 24 TFs (ex; HMBOX1) might target BTK, 71 TFs (ex; BCL11B) might target KIFC2, 61 TFs (ex; TEAD3) might target CALML6, 49 TFs (ex; KAT2A) might target NR4A1, 44 TFs (ex; SP3) might target MYB and 32 TFs (ex; FOSL2) might target ADCY4 and are listed in Table 5.

**Fig. 6.**
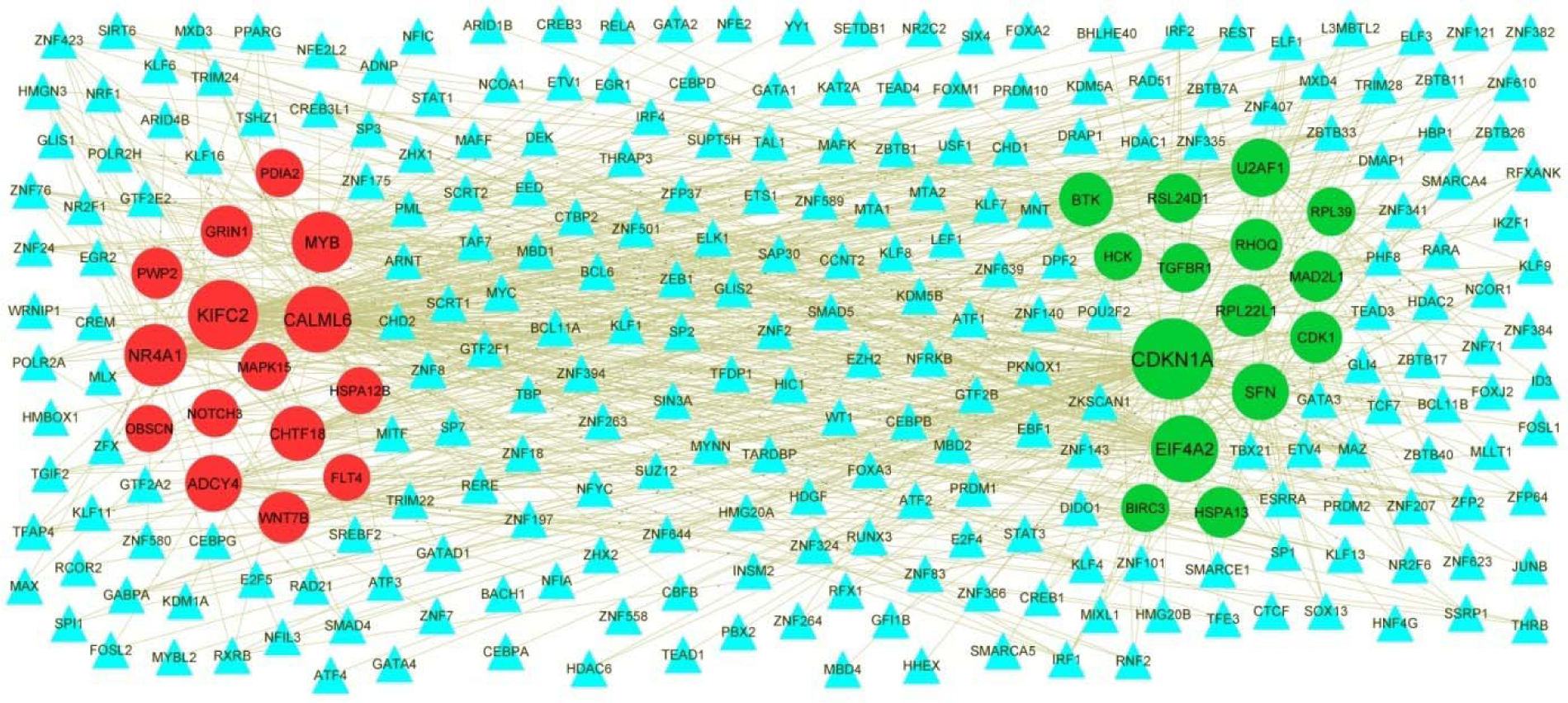
TF - hub gene regulatory network. The blue color triangle nodes represent the key TFs; up regulated genes are marked in green; down regulated genes are marked in red.

### Validation of hub genes by receiver operating characteristic curve (ROC) analysis

ROC curve analysis using “pROC” packages was performed to calculate the capacity of 5 up regulated and 5 down genes to distinguish patients with dementia from healthy controls. CDK1, TOP2A, MAD2L1, RSL24D1, CDKN1A, NOTCH3, MYB, PWP2, WNT7B and HSPA12B all exhibited excellent diagnostic efficiency (AUC > 0.9) (Fig. 7).

**Fig. 7.**
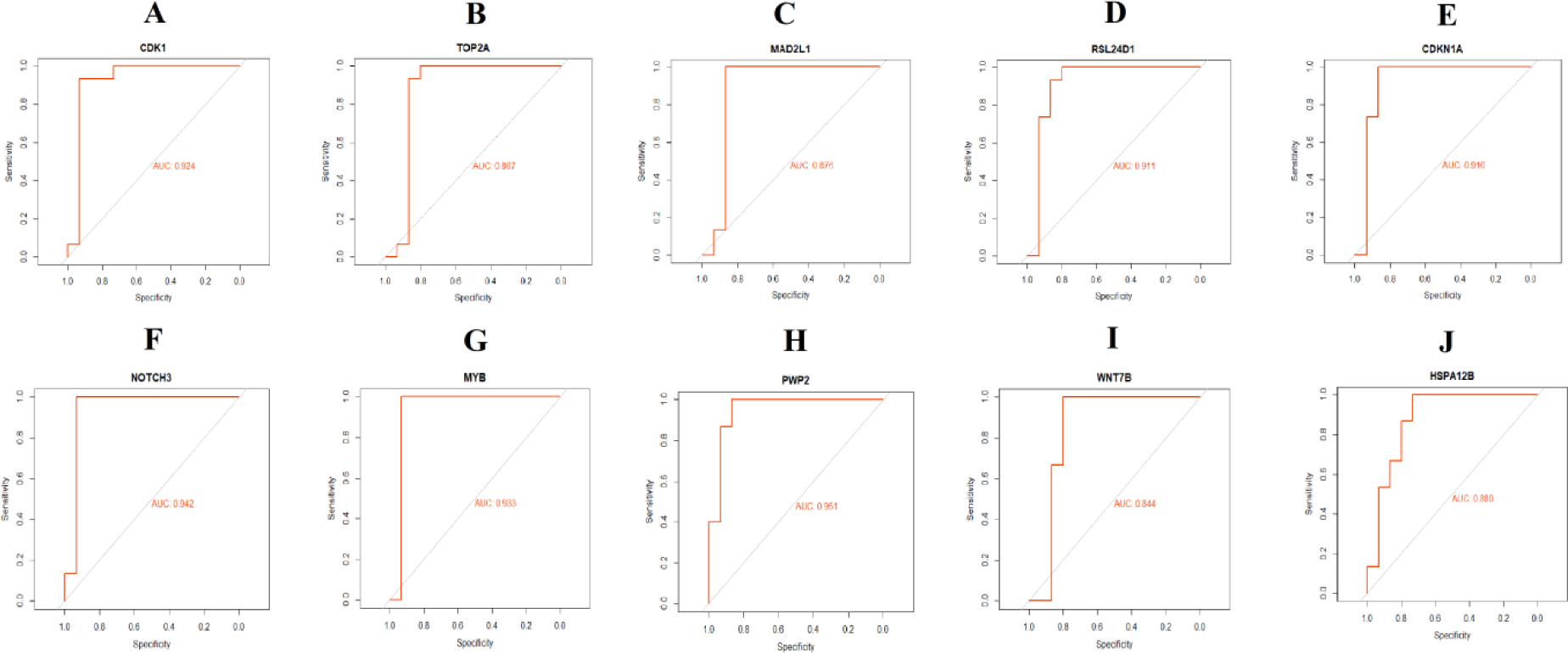
ROC curve validated the sensitivity, specificity of hub genes as a predictive biomarker for dementia prognosis. A) CDK1 B) TOP2A C) MAD2L1 D) RSL24D1 E) CDKN1A F) NOTCH3 G) MYB H) PWP2 I) WNT7B J) HSPA12B

## Discussion

As dementia is the most frequent neurodegenerative disorder worldwide, it is important to understand the molecular mechanism of dementia, which might provide a novel field in diagnosis and therapy of dementia. In this investigation, we got a total of 948 DEGs including 475 up regulated and 473 down regulated genes. [Sanfilippo et al 2020], [Kamboh et al 2006], [Hartlage-Rübsamen et al 2015], [Pey et al 2014], [Yu et al 2015] and [Wang et al 2016] found that CHI3L2, SERPINA3, CCL2, CD163, HLA-DRB5 and GSTM1 are expressed in Alzheimer’s disease patients. BCL2A1 is implicated in the growth of amyotrophic lateral sclerosis [Iaccarino et al. 2011], but this gene might be associated with progression of dementia. HES5 [Kondo et al. 2019] and CYP2D6 [Chamnanphon et al. 2020] are detected in brain cells and is highly correlated with dementia.

The DEGs obtained were remarkably enriched in GO terms and pathway under defense response, whole membrane, molecular transducer activity, neutrophil degranulation, ion transport, integral component of plasma membrane, transporter activity and neuronal system. Defense response [Licastro and Chiappelli, 2003], cell activation [Schubert and Rudolphi, 1998], whole membrane [Paonessa et al. 2019], intrinsic component of plasma membrane [Verdier et al. 2004], neutrophil degranulation [Licastro et al. 1994], innate immune system [van der Willik et al. 2019], hemostasis [Loures et al. 2019], ion transport [Benos et al. 1994], neuron projection [Bondareff et al. 1982], transporter activity [Jellinger et al. 2020], neuronal system [Seeley, 2008] and axon guidance [Zhou et al. 2012] were responsible for progression of dementia. HGF (hepatocyte growth factor) [Sharma, 2010], HLA-DRB1 [Lu et al. 2017], CHI3L1 [Lananna et al. 2020], S100A8 [Lodeiro et al. 2017], S100A9 [Wang et al. 2018], S100A12 [Shepherd et al. 2006], FCGR2B [Costa et al. 2020], HP (haptoglobin) [Song et al. 2015], AIF1 [Sanfilippo et al 2020], CCR1 [Halks-Miller et al. 2003], ALOX5AP [Manev and Manev, 2006], FPR2 [Iribarren et al. 2005], CR1 [Mahmoudi et al. 2018], CFI (complement factor I) [Lashkari et al. 2018], AQP4 [Zeppenfeld et al.2017], SLC11A1 [Jamieson et al. 2005], CTSS (cathepsin S) [Lemere et al. 1995], CXCR2 [Ryu et al. 2015], IL10 [Babi Leko et al. 2020], IL18 [Scarabino et al. 2020], CYP19A1 [Song et al. 2019], BTK (Bruton tyrosine kinase) [Keaney et al. 2019], C7 [Zhang et al. 2019], PLA2G7 [Koshy et al. 2010], SOCS3 [Cao et al. 2018], OLR1 [Wang et al. 2018], LIPA (lipase A, lysosomal acid type) [von Trotha et al. 2006], CD86 [Busse et al. 2015], TLR5 [Herrera-Rivero et al. 2019], CD44 [Pinner et al. 2017], CD68 [Matsumura et al. 2015], CD74 [Kiyota et al. 2015], LTF (lactotransferrin) [Wang et al. 2010], LYZ (lysozyme) [Sandin et al. 2016], S100A10 [King et al. 2020], AQP1 [Nikpour et al. 2020], GPR84 [Audoy- Rémus et al. 2015], HPSE (heparanase) [García et al. 2017], NGFR (nerve growth factor receptor) [Cheng et al. 2012], RHOQ (ras homolog family member Q) [Tian et al. 2021], EGF (epidermal growth factor) [Birecree et al. 1988], CDK1 [Hilgeroth et al. 2014], PKD1 [Chen et al. 2020], HTR6 [Kan et al. 2004], GRIN1 [Wang et al. 2021], AGRN (agrin) [Rauch et al. 2011] and GP6 [Laske et al. 2008] have been shown as a promising biomarkers in Alzheimer’s disease. [Field et al 2010], [Lincoln et al 2009], [Ziliotto et al 2018], El [Sharkawi et al 2019], [Cossins et al 1997], [Mirowska-Guzel et al 2011], [Asouri et al 2020], [Cardamone et al 2018], [Begovich et al 2005], [Bennetts et al 1999], [Iparraguirre et al 2017] and [Oldoni et al 2020] reported that HLA-DPB1, HLA-DQA1, CCL18, CCL20, MMP7, IL1A, IL2RA, CYBB (cytochrome b-245 beta chain), PTPN22, HLA-DMB, ANXA2 and CHIT1expression were associated with progression of multiplesclerosis, but these genes might be liable for advancement of dementia. [Chang et al 2015], [Jamshidi et al 2014], [Campolo et al 2020], [Bottero et al 2018], [Gao et al 2019], [Haga et al 2020], [Klaver et al 2018], [Greenbaum et al 2013] and [Aguirre et al 2020] reported that expression of HLA-DQB1, HLA-DRA, TLR7, TLR8, PTPRC (protein tyrosine phosphatase receptor type C), OSMR (oncostatin M receptor), FABP5, LAMP2, CHRNA5 and IL13RA1 could be an index for Parkinson’s disease progression, but these genes might be responsible for progression of dementia. [Shmuel-Galia et al. 2017], [Paloneva et al. 2000], [Abu-Rumeileh et al. 2020], [Wojta et al. 2020], [Ries et al. 2016], [Bernstein et al. 2020], Qian and Ke [2020], [Kitic et al. 2014], [Gorevic et al. 1985], [Bonham et al. 2018], [Shi et al. 2017], Maddison and Giorgini [2015], [Wang et al. 2017], [Pase et al. 2020], [Daniele et al. 2015], Hayward and Lee [2014], [Wang et al. 2020], [Kim et al. 2006],[Götz et al. 1998], [Budge et al. 2018], [Loeffler et al. 1996], [Cacciagli et al. 2014], [Stoffel et al. 2018], [Koper et al. 2018], [Philippe et al. 2015], [Cursano et al. 2020], [Butler et al. 2019], [Cho et al. 2021] and [Casiglia et al. 2013] presented that expression of TLR6, TYROBP (transmembrane immune signaling adaptor TYROBP), SERPINA1, CCR5, ANXA1, SLC7A2, HAMP (hepcidin antimicrobial peptide), TSLP (thymic stromal lymphopoietin), B2M, CXCR4, C3, KYNU (kynureninase), CASP1, CD14, TLR1, TLR2, TREM2, HCK (HCK proto-oncogene, Src family tyrosine kinase), MAOB (monoamine oxidase B), GPNMB (glycoprotein nmb), CP (ceruloplasmin), AP1S2, SMPD3, CXCL11, SHANK3, CRHR1, DRD4, NOTCH3 and GNB3 were associated with progression of dementia. [Yamamoto et al. 2017], [Wang et al. 2017], [Stringer et al. 2020] and [Brenner et al. 2019] demonstrated that that the expression of SPP1, C5AR1, CACNA1H and CACNA1A are associated with the prognosis of amyotrophic lateral sclerosis, but these genes might be associated with advancement of dementia. TGFBR1 [Zheng et al. 2021], SCN4A [Bergareche et al. 2015], KCNT1 [McTague et al. 2018], SLC13A5 [Matricardi et al. 2020] and CACNA1B [Gorman et al. 2019] were reported to be associated with progression of epilepsy, but these genes might be involved in dementia. A PPI network and modules were constructed with DEGs, which revealed that the top hub genes with highest node degree, betweenness centrality, stress centrality and closeness centrality.TOP2A, MAD2L1, RSL24D1, CDKN1A, PWP2, WNT7B, HSPA12B, CCNG1, GADD45A, SFN (stratifin), CEP55, PTTG1, GRM2, SHANK1 and GRM4 were might be the novel biomarkers for dementia progression.

A miRNA-hub gene regulatory network and TF-hub gene regulatory network with the hub genes were constructed, and identified as the key genes, miRNAs and TFs in dementia. Hsa-mir-1248 [Hewel et al. 2019], hsa-mir- 195-5p [Takousis et al. 2019] and SP3 [Boutillier et al. 2007] might play important role in Alzheimer’s disease, but these genes might be linked with progression of dementia. hsa-mir-101-5p has been reported in multiple sclerosis [Chen et al. 2017], but this gene might be liable for progression of dementia. Reports indicate that hsa-mir-206 [Grasso et al. 2019], hsa-mir-107 [Nelson et al. 2018], STAT1 [Ebner et al. 2011], TBP (TATA-binding protein) [Bruni et al. 2004] and KAT2A [Nematollahi et al. 2016] were found in dementia. BCL11B has been shown to be activated in amyotrophic lateral sclerosis [Lennon et al. 2016], but this gene might be liable for advancement of dementia. [Fan et al. 2020] and [Rouillard et al. 2018] found that FOSL2 and NR4A1 might be involved in the development of Parkinson’s disease, but these genes might be linked with dementia progression. Hsa-mir-6887-3p, hsa-mir-488-5p, hsa-mir-548l, hsa-mir-6886-5p, hsa-mir-150-3p, MLLT1, ELF1, HMBOX1, TEAD3, EIF4A2, HSPA13, KIFC2, U2AF1, CALML6 and ADCY4 were might be the novel biomarkers for dementia progression.

In conclusion, we used a series of bioinformatics analysis methods to identify the key genes and pathways associated in dementia initiation and progression from expression profiling by high throughput sequencing data containing patients with dementia and healthy controls. Our results provide a more detailed molecular mechanism for the advancement of dementia, shedding light on the potential biomarkers and therapeutic targets. However, the interacting molecular mechanism and function of genes need to be confirmed in more experiments.

## Acknowledgement

I thank Delphine Fagegaltier, New York Genome Center, Center for Genomics of Neurodegenerative Disease, Hemali Phatnani, New York, USA, very much, the author who deposited their profiling by high throughput sequencing dataset GSE153960, into the public GEO database.

## Ethical approval

This article does not contain any studies with human participants or animals performed by any of the authors.

## Informed consent

No informed consent because this study does not contain human or animals participants.

## Availability of data and materials

The datasets supporting the conclusions of this article are available in the GEO (Gene Expression Omnibus) (https://www.ncbi.nlm.nih.gov/geo/) repository. [(GSE153960) (https://www.ncbi.nlm.nih.gov/geo/query/acc.cgi?acc=GSE153960)]

## Consent for publication

Not applicable.

## Author Contributions

B. V. - Writing original draft, and review and editing

C. V. - Software and investigation

## Authors

Basavaraj Vastrad ORCID ID: <u>0000-0003-2202-7637</u> Chanabasayya Vastrad ORCID ID: <u>0000-0003-3615-4450</u>

